# Cell-Surface RNA Forms Ternary Complex with RNA-Binding Proteins and Heparan Sulfate to Recruit Immune Receptors

**DOI:** 10.1101/2024.07.23.604836

**Authors:** Zeshi Li, Bhagyashree S. Joshi, Hongbo Yin, Ruud H. Wijdeven, Rue A. Koç, Dick Zijlman, Abhijeet Pataskar, Irene Santos-Barriopedro, Hailiang Mei, Wei Wu, Milad Shademan, Filip M. Zawisza, Eric Bos, Pradeep Chopra, Marvin Tanenbaum, Thomas Sharp, Michiel Vermeulen, Vered Raz, Chirlmin Joo

## Abstract

Recent discoveries have shown the presence of RNA molecules on the cell surface, defying the traditional view that RNA only functions intracellularly. However, it is not well understood how cell-surface RNA (csRNA) is stably present on the plasma membrane and what functions it performs. We answer the pressing questions in the emerging field by taking integrated omic-wide approaches and multiple orthogonal validatory methods. Firstly, we exploited the RNA-sensing ability of TLR7 as a specific recombinant probe to detect csRNA. Coupling it with a genome-wide CRISPR-Cas9-knockout screening, we identified heparan sulfate (HS) as a crucial factor for RNA presentation on cells. Using the TLR7 probe, cell surface proximity labelling revealed that these HS-associated csRNAs (hepRNAs) are in vicinity with a plethora of RNA-binding proteins. The compelling observation led us to a molecular model where HS, RNA and RBP form ternary complexes at cell surface. A photochemical RNA-protein crosslinking technology termed SCOOPS were then established to validate the termolecular model in a TLR7-orthogonal manner. Moreover, enabled by SCOOPS, we unveiled identities of hepRNA using next-generation sequencing, and identified traits in RNA primary structures that facilitate HS association. We further show that hepRNA binds to killer cell immunoglobulin-like receptor 2DL5 (KIR2DL5), recruiting the protein to cell surface and potentially enhancing receptor-ligand interactions. Our findings provide a foundation for exploring how cell-surface ribonucleoproteins contribute to immune modulation.

## Introduction

Ribonucleic acid (RNA) performs many critical cellular functions in both prokaryote and eukaryote cells. Although the localization of RNA was considered primarily intracellular in mammalian cells, emerging studies have indicated that RNA can be displayed on the cell surface. A previous study found that bacterial genome-encoded RNA can be anchored on their cell surface via binding to a protein.^1^ Very recent studies have shown the presence of RNA on the extracellular part of mammalian cell plasma membranes.^2–5^ These cell-surface-associated RNAs (csRNAs) include fragments of messenger and long non-coding RNAs and putative glycosylated small RNAs. There are indications that csRNAs can participate in molecular recognition, thereby modulating cell-cell interactions. A recent study has shown csRNA mediated neutrophil recruitment by interacting with P-selectin on the endothelial cells.^6^ Despite the aforementioned studies, mechanistic understanding as to how RNAs are stably presented on the mammalian cell surface remains limited. It is also unclear if csRNA may exhibit other modes of function besides presenting glycan moieties to interacting cells.^2,6^

As most RNA molecules are associated with RNA-binding proteins (RBPs), it can be speculated that csRNA is similarly associated with RBPs, providing a mechanism for cell surface anchoring. In line with this hypothesis, both earlier and recent studies have demonstrated RBPs, previously thought to be localized only intracellularly, are also present on the plasma membrane.^7–10^ Thus, the potential binding partners are indeed available for and in proximity to RNA on the cell surface. However, these RBPs are generally soluble proteins that do not possess a transmembrane domain and are therefore unlikely to be directly anchored on the lipid bilayer like many cell-surface glycoproteins. For example, a classical nuclear RBP, U5 SNRNP200, has been evaluated as an acute myeloid lymphoma-associated cell surface antigen for immunotherapy.^9, 10^ This RBP was shown to be associated with the Fcγ receptor IIIA. A more recent study revealed that a plethora of RBPs can localize to the cell surface, as possible carriers of csRNA.^11^ However, it is largely unknown how csRNA and RBPs become stably presented on the plasma membrane, and if so, whether they are associated with each other.

Animal cells employ highly negatively charged linear carbohydrate polymers, such as glycosaminoglycans, as a scaffold to enrich soluble proteins and organize membrane bound proteins on the plasma membrane.^12^ Heparan sulfate (HS) is an important class of glycosaminoglycans ubiquitously found on plasma membrane and in extracellular matrix (ECM).^13^ Portions of HS are heavily sulfated, and these sulfated domains serve as docking sites for a broad array of proteins, including growth factors, morphogens, amyloid proteins, lipoproteins, cytokines, and chemokines, as well as many microbial proteins. HS is characterized by having an extension of repeating disaccharide units, glucosamine-α1,4-glucuronic acid (GlcN-GlcA) or glucosamine-α1,4-iduronic acid (GlcN-IdoA). An HS chain can contain over a hundred monosaccharide units. Glycan modifications such as N- and O-sulfation introduce negative charges to portions of the HS chain and are known to be crucial for the biological functions of HS. The pattern of N- and O-sulfation, also known as the sulfation code, critically determines what proteins interact with HS. These features make HS among a main director of cell surface interactions.^14^

Herein we demonstrate that HS functions as a scaffold for presenting RNAs and RBPs on the cell surface. A key tool in our study was toll-like receptor 7 (TLR7), which we leveraged as a probe for csRNA. We established that the recombinant TLR7-Fc fusion protein can effectively detect and capture RNA on living cells, enabling a genome-wide knockout screening to identify genes regulating csRNA localization. The screening revealed HS biosynthetic enzymes as essential factors. Using the same probe, we also identified the csRNA-proximal proteome, which includes both classical and non-canonical RBPs. These findings led us to propose a model in which RNA, RBP, and HS form ternary complexes for csRNA presentation. To further validate this ternary configuration, we developed an TLR7-independent approach—Spatio-selective Crosslinking followed by Orthogonal Organic Phase Separation (SCOOPS). SCOOPS employs local photocatalytic generation of singlet oxygen (SO) to crosslink RNA with bound proteins, facilitating selective isolation of csRNA. By combining SCOOPS with next-generation sequencing, we identified HS-associated csRNAs and uncovered primary structural imprints contributing to their localization. Finally, we demonstrated that HS-associated csRNAs can recruit killer cell immunoglobulin-like receptor 2DL5 (KIR2DL5) to the cell surface, suggesting a potential role for csRNA in modulating immune interactions between KIR2DL5 on immune cells and its cognate ligand on target cells.

## Results

### Establishment of TLR7-Fc as a probe for csRNA

The sequence of RNA molecules on the plasma membrane likely varies between different cells, and RNA molecules present at low copy numbers may elude detection.^15^ The state-of-the-art csRNA labeling approaches either require fixation or rupturing of the plasma membrane^3, 15^ and are therefore not compatible with live cells. To overcome this, we reasoned that the use of an RNA-recognizing probe with low sequence specificity, rather than a hybridizing probe which requires prior knowledge of csRNA sequence, should boost the chance for csRNA detection, and yield strong signals. We, therefore, exploited nature’s RNA-sensing machinery: toll-like receptor 7 (TLR7),^16^ an endosome-localized, pattern recognition receptor in select immune cells to sense both foreign and endogenous single-stranded RNA molecules.^17–19^ TLR7 requires two consecutive uridine moieties for strong RNA binding, but requires only few other sequence features.^20^ We used the ectodomain of TLR7 (Ala27-Asn839) fused to the human Fc tag (denoted hereafter TLR7-Fc). The former is expected to recognize csRNA on intact (live) cells, and the latter will allow for the detection by a secondary antibody for different downstream applications.

We employed confocal microscopy to examine the efficiency of TLR7-Fc to detect csRNA. TLR7-Fc is applied onto intact, fixed cells without cell permeabilization, followed by washing and the addition of a fluorophore-conjugated, anti-human IgG secondary antibody. To check for localization of TLR7-Fc staining, we labeled the cell surface with biotin-conjugated wheat germ agglutinin (WGA-biotin),^21^ a lectin that recognizes N-acetylneuraminic acid and N-acetylglucosamine residues present in cell surface glycans, and the biotin moieties were detected by fluorescent streptavidin. We observed that bright puncta given rise to by TLR7-Fc probe in complex with the fluorescent secondary antibody align with the rims of HeLa, U2OS, and Mel526 cells (**Figure 1a** and **Supplementary Figure 1a** and **b**). This is akin to many cell surface proteins, which manifest in clusters.^22, 23^ WGA signal remained confined to the cell surface. csRNA signal was absent in controls devoid of TLR7-Fc but only including the secondary antibody (**Supplementary Figure 1c**). A treatment with RNaseA/T1 cocktail in fixed, non-permeabilized cells, severely disrupted the TLR7 signal (**Figure 1a**), while DNase treatment did not (**Supplementary Figure 1d**), validating the RNA specificity of TLR7 binding. Moreover, the pre-saturation of TLR7-Fc with an exogenous, synthetic single-stranded RNA competitively inhibited TLR7 binding, further supporting the specificity of TLR7 to probe RNA on cells (**Figure 1a**).

**Figure 1.**
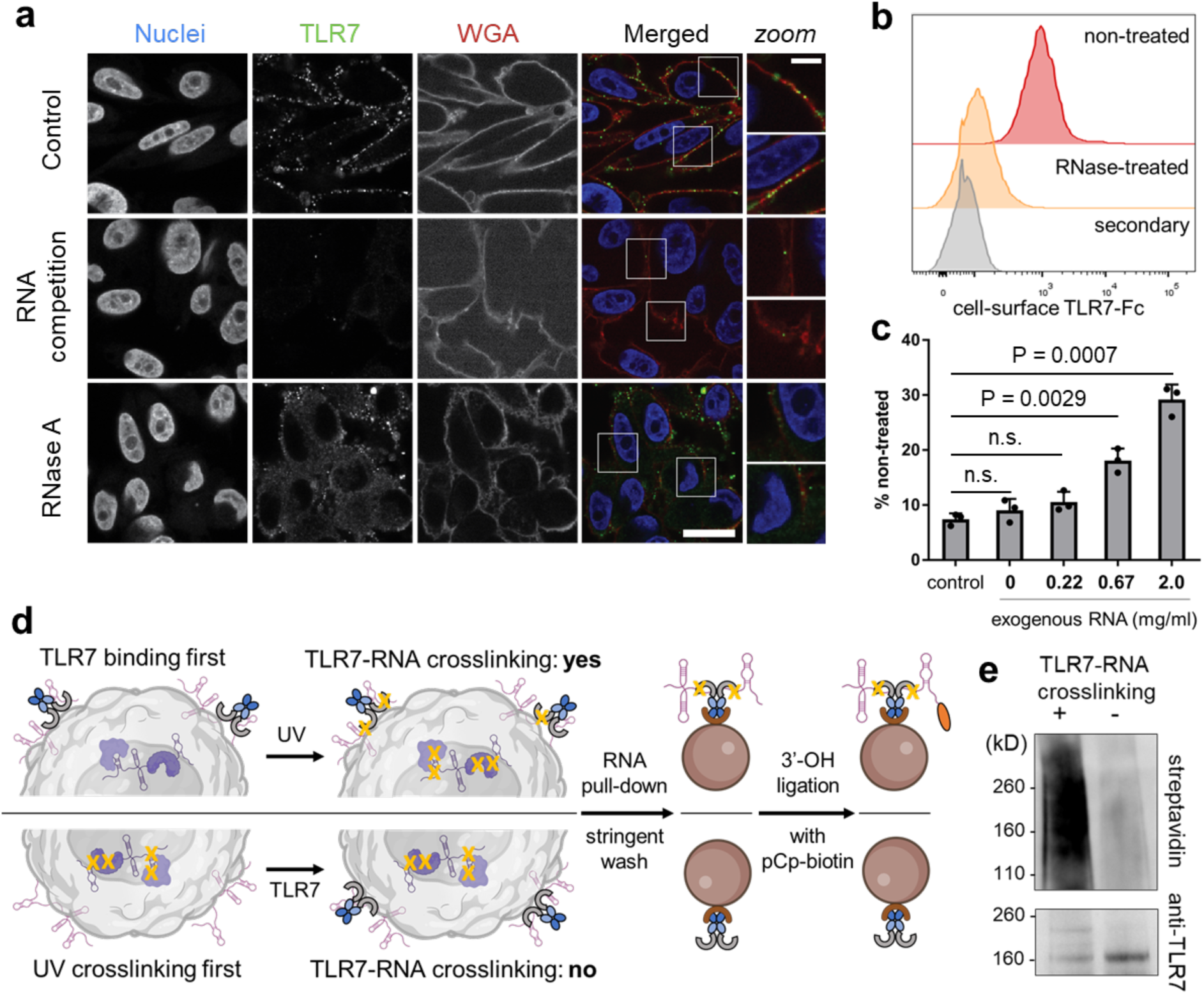
TLR7-Fc is a csRNA probe. **a**) Confocal microscopy images of csRNA on Mel526 cells probed by TLR7-Fc and AlexaFluor488-conjugated Goat-anti-human IgG. Cell surface was stained by WGA-biotin and streptavidin-AlexaFluor594. See supplementary information for the sequence of competitor ssRNA. RNase A concentration, 40 μg/ml. Nuclear counterstaining: DAPI. Scale bar: 20μm. **b**) Histograms from flow cytometry analysis of csRNA on Mel526 cells probed by TLR-Fc in complex with goat-anti-human-AlexaFluor647. 40 μg/ml RNase A was used to deplete csRNA during EDTA lifting. n = 3 independent cell cultures. **c**) TLR7-Fc binding was rescued by exogenous total RNA in a concentration-dependent manner. n = 3 independent cell cultures. Exogenous RNA in 1x DPBS was added to cells at 4 ℃. 0 mg/ml RNA means 1x DPBS buffer without RNA. The control was with cells treated with RNase A but without any additional incubation. Y-axis shows relative signal intensity, with the geometric mean of TLR7 binding on non-RNase treated cells set to 100%. Two-tailed Student’s t-test, mean ± s.d. **d**) Schematics of in-situ crosslinking and csRNA capture experiment. pCp, cytidine-5’-phosphate-3’- (6-aminohexyl)phosphate. Orange oval represents biotin. **e**) Chemiluminescent detection of TLR7-captured and biotin-labeled RNA. Rabbit-anti-TLR7 was used to check for TLR7-Fc on the same nitrocellulose membrane after blotting biotinylated RNA. n = 3 independent cell cultures for crosslinked and non-crosslinked samples.

Live-cell staining with TLR7 showed the same puncta pattern, indicating that the clustering of our probe on the cell surface is unlikely to be caused by cell fixation (**Supplementary Figure 1e**).^24–26^ Interestingly, the monoclonal antibody J2, previously used for intracellular detection of endogenous dsRNA^27^ and viral dsRNA,^28^ as well as for cell-surface glycoRNA,^2^ did not give detectable signals on HeLa or U2OS cell surface (**Supplementary Figure 1f**). Next, to observe csRNA in a more detailed subcellular context, we employed electron microscopy (EM), leveraging the TLR7-Fc probe and Protein-A-gold (10 nm) as a secondary probe (**Supplementary Figure 1g** and **h**). We observed csRNA signals along the cellular periphery, cell-cell junctions, and endocytosed vesicles, confirming the findings with fluorescent microscopy.

We further used our TLR7-Fc probe to quantify csRNA and its sensitivity to RNase treatment with flow cytometry (**Figure 1b** and **Supplementary Figure 2a**). Corroborating the fluorescence imaging results, when live cells were pre-treated with RNase A, the signal intensity given rise by TLR7-Fc in complex with the fluorescent secondary antibody dropped dramatically close to the background. To further confirm that TLR7-Fc binds to RNA on the cell surface, the prior csRNA-depleted cells by RNase were incubated with exogenous, purified total cellular RNA (**Supplementary Figure 2b**). We found that TLR7 binding could be partly restored by exogenous RNA in a concentration-dependent manner (**Figure 1c**).

To further validate that TLR7 binds to RNA on the cell surface, we performed in-situ RNA-protein crosslinking with UVC after TLR7-Fc probe addition, followed by RNA precipitation and RNA 3’-end labeling (**Figure 1d**). Upon UVC irradiation, csRNA is expected to be covalently bound to the TLR7-Fc probe and can be co-precipitated with the probe using protein A-functionalized magnetic beads. The precipitated RNA was then labeled at the 3’-end with biotin-cytidine using RNA ligase 1. This was followed by SDS-PAGE and Western blot for biotin. As a negative control, cells were UVC irradiated before TLR7-Fc incubation and subjected to the same procedure. UVC-crosslinked biotinylated csRNA is expected to remain bound to TLR7 throughout the entire procedure, whereas the csRNA not crosslinked to TLR7 will dissociate upon denaturation, and therefore will not remain on the membrane.^29^ We observed a strong chemiluminescent signal above TLR7-Fc molecular weight for crosslinked csRNA-TLR7 (**Figure 1e**), but not for non-crosslinked TLR7 probe. The results demonstrate TLR7 probe indeed binds to RNA on the cell surface.

To confirm TLR7-Fc binding to RNA is mediated by TLR7 ectodomain, but not other portions of the protein (such as the Fc tag), we performed an in-vitro RNA crosslinking and co-precipitation assay using the TLR7-Fc and human IgG control. Fragmented, purified total cellular RNA was incubated and crosslinked (irradiated with UVC, 254 nm) with TLR7-Fc and IgG, pulled down on beads and intensively washed, labeled with Cy5 at 3’-end, and finally released from beads by proteinase K digestion. Purified precipitated RNA were then analyzed in agarose gel electrophoresis. Strong fluorescent bands were observed for TLR7-Fc captured RNA, but not for IgG control (**Supplementary Figure 2c**). The results indicated the Fc tag did not non-specifically bind to RNA, demonstrating the RNA specificity of our probe.

### Genome-wide screening identified essential factors for csRNA presentation

Previous studies have only postulated that csRNA can be presented on the plasma membrane, yet a detailed mechanistic understanding of this process is lacking. With TLR7-Fc as a general csRNA probe in hand, we next sought to uncover the molecular underpinnings of csRNA presentation. We performed a pooled genome-wide, CRISPR-Cas9-mediated knockout (KO) screening^30^ to identify essential cellular factors for csRNA stable presentation. TLR7-Fc in complex with a fluorescent secondary antibody was used to enrich the csRNA^low^ population with fluorescence-assisted cell sorting (FACS). Such FACS-based enrichment was performed twice. The enriched cells together with the non-enriched input cells were subjected to deep sequencing to identify the guide RNAs of CRISPR-Cas9 giving rise to the csRNA^low^ phenotype. (**Figure 2a**, see **Supplementary Figure 3** for gating strategies). Likewise, the csRNA^high^ phenotype was also enriched and sequenced (See **Supplementary Text 1** for details, and **Supplementary Table 1** for full results).

**Figure 2.**
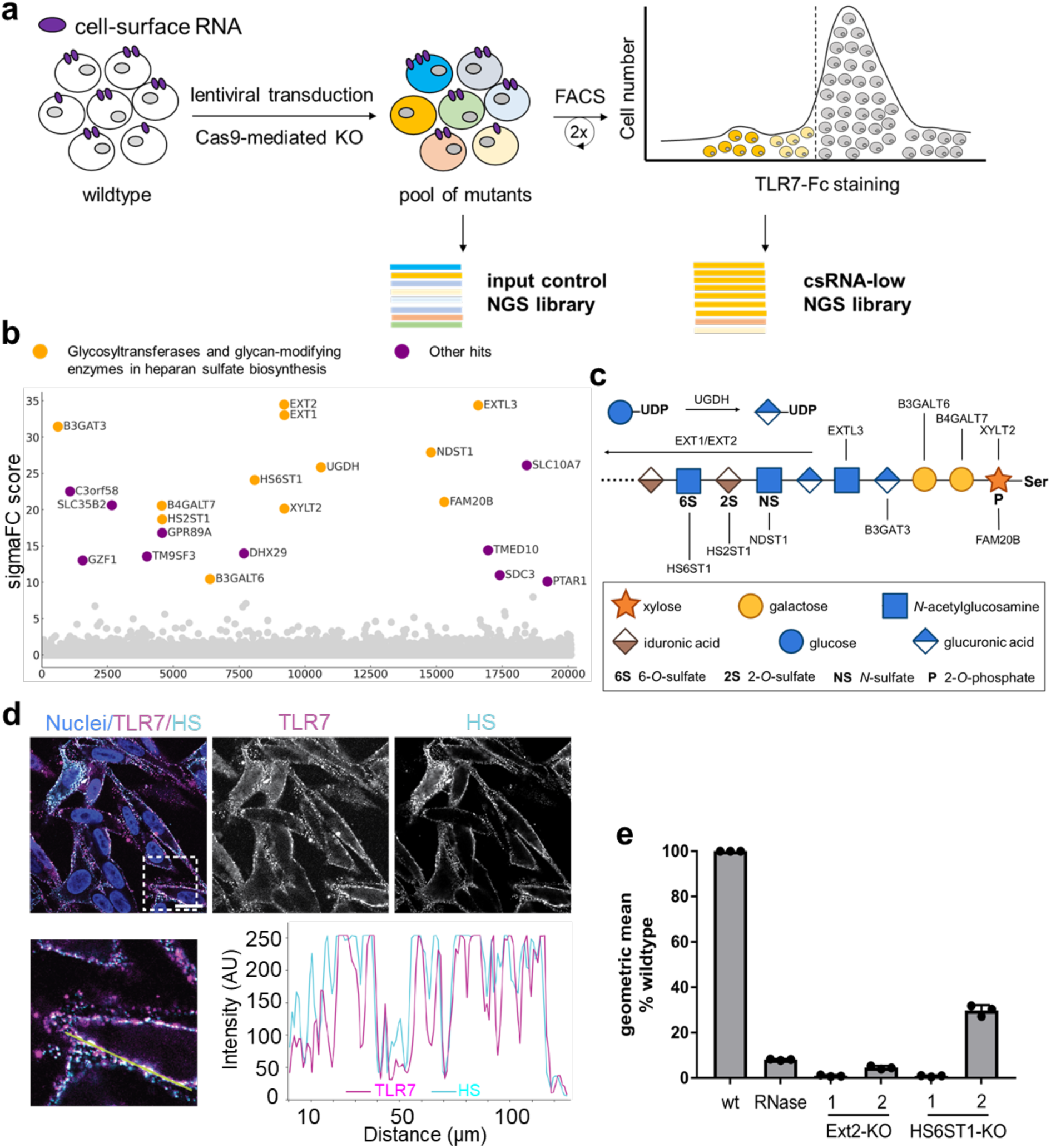
Genome-wide screening reveals essential factors for csRNA stable presentation. **a**) Schematics of the genetic screen in Mel526 cells with csRNA stained with TLR7-Fc and AlexaFluor488-conjugated goat-anti-human IgG. Color-coded cells represent the 5% of cells with the lowest fluorescent signal, indicating the loss of csRNA. **b**) Dot plot of genetic screening results. Y-axis in this plot is the geometric mean of the SigmaFC scores from the duplicates calculated using PinAPL-Py. Heparan sulfate-related glycosyltransferases and glycan-modifying enzymes are colored in yellow. Other hits with a SigmaFC larger than 10 are colored in purple. **c**) HS biosynthetic pathway. Gene candidates as yellow dots in Panel **b** are shown in this schematic representation. **d**) csRNA and HS colocalization on cell surface of Mel526. csRNA was probed by TLR7-Fc and AlexaFluor647-conjugated rat-anti-human IgG (magenta), while HS was labeled with mouse anti-HS (10E4 epitope) antibody and AlexaFluor488-conjugated goat anti-mouse IgM mu chain secondary antibody (cyan). The line intensity profile along the yellow line shows csRNA tightly overlaps with heparan sulfate. Nuclear staining (blue): DAPI. Scale bar: 20 μm. The frame with dashed line indicates the area where intensity profile was collected. **e**) Flow cytometry analysis showing csRNA loss in EXT2 and HS6ST1 KO mutant Mel526 cells. TLR7-Fc with goat-anti-human-AlexaFluor647 was used for csRNA staining. 40 μg/ml RNase A treated cells are used as control. The numbers above EXT2 and HS6ST1 represents different guide RNAs for KO (see Materials and Methods for sequences). n = 3 independent cell cultures.

Data analysis revealed that enzymes involved in heparan sulfate (HS) biosynthesis^31^ were among the most highly scored hits, suggesting HS is a pivotal molecular factor for the stable cell-surface display of RNAs (**Figure 2b**, full list of hits in **Supplementary Table 2**). Our hits include enzymes for the core tetrasaccharide synthesis (XYLT2, B4GALT7, B3GALT6, and B3GAT3), glycan chain polymerizing enzymes (EXT1, EXT2, and EXTL3), as well as glycan chain modifying enzymes such as sulfotransferases (NDST1, HS2ST1, and HS6ST1) and a xylose kinase (FAM20B) (**Figure 2c**). These hits suggest the stable presentation of csRNA requires a fully extended HS chain and appropriate N- and O-sulfation pattern.

Apart from the HS-related glycosyltransferases and glycan-modifying enzymes, we also identified regulators in HS biosynthesis. For example, UGDH oxidizes uridine diphosophate linked glucose (UDP-Glc) to UDP-GlcA, a glycosyl donor for HS polymerization; SLC35B2 is a Golgi membrane transporter of the sulfo-donor, adenosine 3’-phospho-5’-phosphosulfate;^32^ SLC10A7 is crucial for cellular calcium regulation and was implicated in HS biosynthesis;^33^ and C3orf58 has been recently shown to regulate HS biosynthesis.^34^ The genes related to the N-glycan biosynthetic pathway were not among the candidates, suggesting the csRNA detected by TLR7 is independent from glycoRNA, and it does not require N-glycosylation to access plasma membrane.

### Validation of genome-wide screening

Next, our attention was focused on validating that HS is necessary for a display of the csRNA detected by the TLR7-Fc probe. csRNA location closely correlated with the heparan sulfate staining on the cell surface showing their tight proximity (**Figure 2d**). We then generated HS-deficient mutant cells lacking either the elongation enzyme EXT2 or sulfotransferase HS6ST1 via CRISPR-Cas9 mediated KO, each with two high-scoring guide RNAs, and used flow cytometry to quantify TLR7-Fc binding. Both guide RNAs for EXT2 resulted in an almost complete loss of TLR7 binding. One guide RNA for HS6ST1 led to a similar phenotype as did EXT2 KO, whereas the other guide RNA reduced TLR7 binding down to approximately 30% of the wildtype (**Figure 2e**). In consistency with the flow cytometry experiments, confocal microscopy also showed a dramatic reduction in csRNA signal around the cell rim after knocking out EXT2, HS6ST1, EXTL3, and C3orf58 (**Supplementary Figure 4a**). The loss of TLR7 binding to the cell rim was also observed in Chinese hamster ovary (CHO) cell mutants (pgsD-677 and pgsE-606) deficient in HS biosynthesis (**Supplementary Figure 4b**). The wildtype CHO cell showed a similar punctate pattern as did Mel526.

An alternative scenario which might have given rise to similar hits in our genome-wide screening would be that TLR7-Fc directly binds to HS on cell surface (**Supplementary Figure 4c**). Thus, the loss of TLR7 binding to cell surface would have been caused by the RNase non-enzymatically and competitively suppressing TLR7-Fc binding to HS. To exclude this, we compared the TLR7-Fc to well-studied HS-binding proteins using flow cytometry, including fibroblast growth factor-2 (FGF2) and an anti-HS IgM 10E4.^35^ All proteins bound to wildtype cell surface strongly, but hardly to EXT2-KO cells, confirming the HS dependency of the tested proteins (**Supplementary Figure 4d**). However, the two HS binders differed from TLR7 by that they were refractory towards RNase treatment. In addition, we performed surface plasmon resonance binding studies of the RNase (Purelink RNase A) used in the above studies with heparin-coated chips (**Supplementary Figure 4e**). No signal was detected across all RNase concentrations, suggesting the RNase used in the study did not bind to HS, therefore unlikely to competitively suppress other proteins for HS binding. These results strengthened TLR7 as a csRNA probe and demonstrated its loss of binding to cell surface was likely due to the RNase-mediated digestion of csRNA.

To further validate the genetic screen results, we selected two HS-deficient mutant cells for TLR7 binding rescue experiment by exogenous total RNA. Cells were first treated with RNase A, and then with total RNA fragments at varying concentrations (**Supplementary Figure 4f**). The treated cells were then incubated with TLR7-Fc followed by a fluorescent secondary antibody. The TLR7 binding remained close to the background throughout all experimental groups regardless of the RNA concentrations. These results demonstrated that cells lacking an extended HS chain or an appropriate sulfation pattern become refractory to exogenous RNA incubation and can no longer rescue TLR7 binding, suggesting that csRNA presentation requires intact HS. Hereafter, we refer the csRNA detected by the TLR7-Fc probe as heparan sulfate-associated RNA (hepRNA).

### RNA, RBP and HS are in vicinity on cell surface

Both RNA and HS are highly negatively charged biopolymers. How are these two macromolecules associated? We were struck by the observation that upon proteolytic detachment of adherent cells by TryPLE before flow cytometry, TLR7-Fc no longer detectably bound to the cell surface (**Supplementary Figure 5a**), while non-tryptic detachment methods such as EDTA allowed for hepRNA detection by TLR7-Fc (**Figure 1b**). The observation led us to hypothesize that hepRNA may require an additional factor, such as proteins, to be presented on the cell surface HS (**Supplementary Figure 5b**). We, therefore, applied a peroxidase-mediated proximity labeling approach^36^ to identify the hepRNA-proximal proteome using TLR7-Fc. Peroxidase-mediated proximity labeling has been shown to exhibit particularly high efficiency within the radius of ∼20 nm due to the direct contact of peroxidase catalytic center with proximal proteins. Decreased, but considerable labeling does occur within a radius of ∼270 nm,^37, 38^ due to the diffusion of peroxidase-activated biotin-phenol radical.^38^ Thus, peroxidase-based proximity labeling is expected to afford protein candidates that are direct binders of hepRNA, and those that do not physically interact with hepRNA, but are spatially close to it.

We pre-complexed TLR7-Fc with horse radish peroxidase (HRP)-conjugated protein A, which was then incubated with untreated and RNase A-pretreated Mel526 cells (**Figure 3a**). RNase treatment is expected to digest unprotected regions of all csRNA, whereas the portions proximal to or bound by a protein are protected from digestion (**Figure 3b**), an effect commonly exploited to study proteins’ footprint on an RNA.^29, 39^ As an isotype control, human IgG was used instead of TLR7-Fc. Biotinylation of proteins on the cell surface was confirmed by fluorescence imaging and Western blot (**Supplementary Figure 5c** and **d**). Following proximity labeling and non-denaturing cell lysis, biotinylated cell-surface proteins were enriched with streptavidin-functionalized agarose resin. Bound proteins were then on-bead digested with trypsin and subjected to LC-MS/MS and label-free quantitative proteomics analysis.

**Figure 3.**
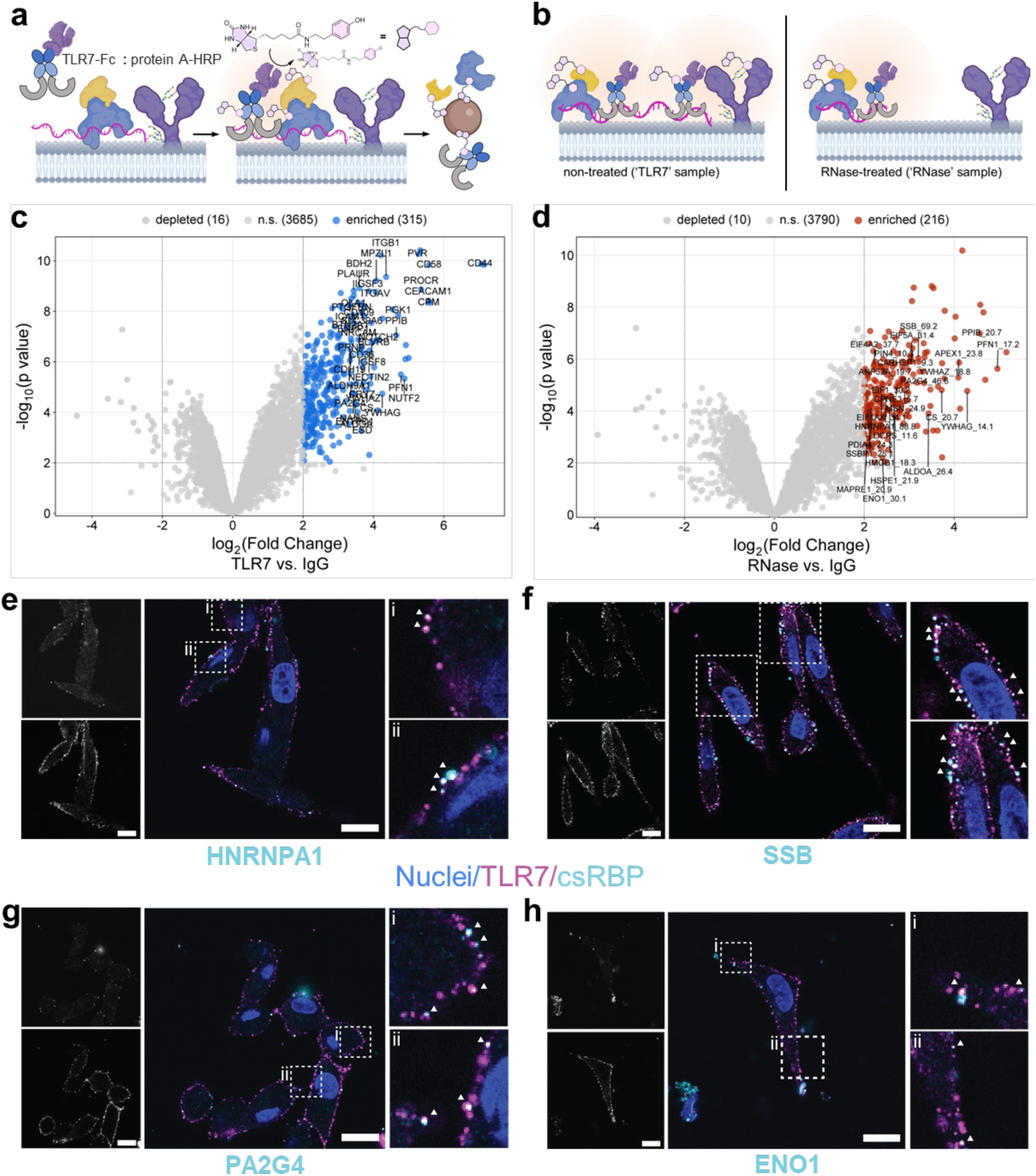
TLR7-proximal proteome harbors known RBPs. **a**) Schematics of TLR7-mediated proximity labeling to identify hepRNA-proximal proteome. As negative control, IgG was used instead of TLR7-Fc for forming complex with protein A-HRP. **b**) Extracellular RNase treated cells were also proximity-labeled in the same fashion. **c**) Volcano plot from the comparison of proteome between TLR7-mediated proximity labeling and IgG control. hepRNA-proximal candidates (fold change ≥ 4, p < 0.01. n = 3 independent cell cultures) are shown in blue. Classically cell-surface localized proteins are labeled with their names. **d**) Volcano plot from the comparison of proteome between TLR7-mediated proximity labeling post-RNase treatment and IgG control. hepRNA-proximal candidates are shown in red. RBP2GO database-documented proteins (RBP2GO score > 10) are labeled with their names and the RBP2GO scores. **e**) – **h**) Confocal microscopy was performed to confirm cell-surface colocalization with hepRNA for select csRBP candidates. The csRBPs were detected with rabbit anti-RBP antibodies and Cy3-conjugated goat anti-rabbit secondary antibody. Nuclear staining: DAPI. Scale bars: 20μm.

By comparing the TLR7-proximal proteome of cells without RNase treatment (denoted as ‘TLR7’) to that of the IgG isotype control, we identified 315 proteins as significantly enriched hits (fold change ≥ 4 AND p value < 0.01. Full list of identified proteins in **Supplementary Table 3**). Of the significantly enriched hits, 137 of them carry the gene ontology (GO) term ‘cell surface’ (GO:0009986) or ‘extracellular region’ (GO:0005576), including known glycoproteins and glycosylphosphatidylinositol-anchored proteins (**Figure 3c**). 76 of the cell-surface (glyco)proteins were not anymore enriched over IgG control in the RNase-treated cell dataset (denoted as ‘RNase’) (**Supplementary Figure 6a**), which suggested upon extracellular RNase digestion, hepRNA is no longer proximal to the (glyco)proteins.

Across the TLR7 and RNase datasets, a considerable number of proteins remained consistently enriched, with a similar fold change (vs. IgG control) in both datasets (**Figure 3d** and **Supplementary Figure 6b**. See full list in **Supplementary Table 4**). In the RNase sample, these proteins likely had been proximity-biotinylated by the residual TLR7-Fc:protein A-HRP complex bound to the RNase-truncated hepRNA (**Supplementary Figure 6c**). We queried RBP2GO, a database documenting validated and putative RBPs with the consistent hits from both TLR7 and RNase samples. It returned with a match of 137 proteins. Among these RBP2GO proteins, 34 carried the GO term ‘RNA binding’ (GO:0003723). The RBP2GO-documented hits include classical RNA-binding proteins such as heterogeneous nuclear ribonucleoprotein A1 and A2B1 (HNRNPA1 and HNRNPA2B1, respectively), and Sjögren syndrome antigen B (SSB), all of which bear a high RBP2GO score, indicating high confidence for direct RNA binding. Non-canonical RNA binding proteins that do not possess a classical RNA-binding domain were also among the hits, many of which are metabolic enzymes or chaperones with moonlighting RNA-binding functions, such as enolase 1 (ENO1)^40^ and calreticulin (CALR).^41^ These proteins normally are scored moderately in RBP2GO database.

We next asked if the detection of the RBPs had been because they were on the cell surface or a result of non-specific labeling/enrichment of intracellular components. We selected both classical and non-canonical RBP hits spanning a wide range of RBP2GO scores for immunofluorescence imaging to check their localization (**Figure 3e** – **h**). All proteins tested, including HNRNPA1, La protein, PA2G4 and ENO1 were found on cell surface, and they were partially colocalizing with TLR7-Fc, suggesting hepRNA and a fraction of these RBPs are in vicinity. Flow cytometry analysis of cell-surface HNRNPA1 and ENO1 (**Supplementary Figure 6d**) with the EXT2-KO mutant cells showed that HS deficiency resulted in a loss of the cell-surface localized RBPs (csRBPs). Western blot of whole cell lysate from wildtype and EXT-KO cells confirmed the absence of csRBP was due to the lack of HS as a scaffold on cell surface, but not a global reduction of protein expression because of HS deficiency (**Supplementary Figure 6e**).

The spatial proximity of csRBP and hepRNA prompted us to examine the formation of ribonucleoprotein complexes (RNPs). Recent years have seen methodological innovation on facile isolation of RNA-bound proteins coupled with MS proteomics to study their dynamics in cells. Several acid guanidinium thiocyanate-phenol-chloroform (AGPC) phase separation strategies^42–44^ leverage the amphiphilicity of UV-crosslinked RNPs for isolation and are independent of RNA sequence. Although UV crosslinking is applied to the whole cell, selectively tagging the surface proteome on intact cells should enable enriching RNA-bound proteins at cell-surface after AGPC phase separation (**Supplementary Figure 6f**). If the RBPs are not bound by RNA, the former should be washed away during the phase separation procedure. We employed a cell-impermeable, amine-reactive biotinylation agent to tag the surface proteome, after which we performed UVC crosslinking and orthogonal organic phase separation (OOPS)^42^ to isolate the crosslinked whole-cell RNPs. The total RNA-bound proteins were then released from the cleaned interphase by RNase digestion and a subsequent APGC extraction into a new organic phase and finally subjected to biotin-based affinity pulldown for label-free quantitative proteomics. A non-biotinylated, but crosslinked sample was taken along to control for non-specific bead binding by the AGPC-denatured proteins.

649 proteins were significantly enriched over the no-biotin control (fold change ≥ 8 AND p value < 0.01) (**Supplementary Table 5**). Enriched GO molecular function terms were heavily related to RNA, demonstrating the effectiveness of the procedure to isolate RBPs (**Supplementary Figure 6h**). Despite the vast difference in the working principles and chemo-selectivity between the TLR7-based proximity labeling and biotinylation-crosslinking-OOPS methods, the latter validated 41 proteins in the TLR7-proximal datasets (TLR7 vs. IgG) as RNA-bound proteins on cell surface (**Supplementary Figure 6g**) in a high-throughput manner. Taken together, the series of proteomics and validatory studies pointed to a molecular configuration of hepRNA, csRBP and HS, in which these three types of biomolecules form ternary complexes.

### HS’s essential role in presenting hepRNA-csRBP complexes

We have demonstrated the RNA’s dependency on and the colocalization with HS, and that these hepRNAs are in the state of RNP complexes. Next, we sought to further validate that the three biomolecules form ternary complexes on cell surface. We introduce herein a facile strategy for cell-surface RNP isolation, termed Spatio-selective Crosslinking followed by OOPS (SCOOPS). SCOOPS allows for one-step isolation of crosslinked RNPs within a subcellular region, without separate tagging and affinity pulldown steps. It is based on our original, unexpected finding that singlet oxygen (SO), a highly reactive oxygen species, can lead to RNA-protein crosslinking (**Supplementary Figure 7**. See **Supplementary Text 2** for a detailed description), an effect analogous to conventional UVC irradiation. However, unlike UVC crosslinking, which can only be applied to whole cells, numerous reports found SO can be generated locally within a subcellular region due to its transient nature (half-life in water ∼4 μs^45^) to achieve spatio-selective chemistry.^46–48^ Therefore, we envisioned that by tethering a SO generator to cell surface, locally produced SO could lead to crosslinking of RNA to bound proteins selectively at cell surface. Subsequently, crosslinked cell-surface RNA-protein complexes were brought into the AGPC interphase for isolation by OOPS.

To establish SCOOPS for cell surface biomolecules, we tethered a small-molecule SO generator, eosin Y (EY) onto streptavidin (strep-EY). Strep-EY retained full binding capacity to biotin (**Figure 4a** and **b**) and can be recruited to live cell surface by WGA-biotin (**Figure 4c**). We expect the AGPC interphase to contain crosslinked csRNA-protein only when csRNA is intact AND when both WGA-biotin and strep-EY are present, while missing out any component would afford little, if any, csRNA after SCOOPS (**Figure 4d**). To perform SCOOPS, cells were incubated sequentially with WGA-biotin and strep-EY and were subjected to AGPC immediately after green light irradiation.

**Figure 4.**
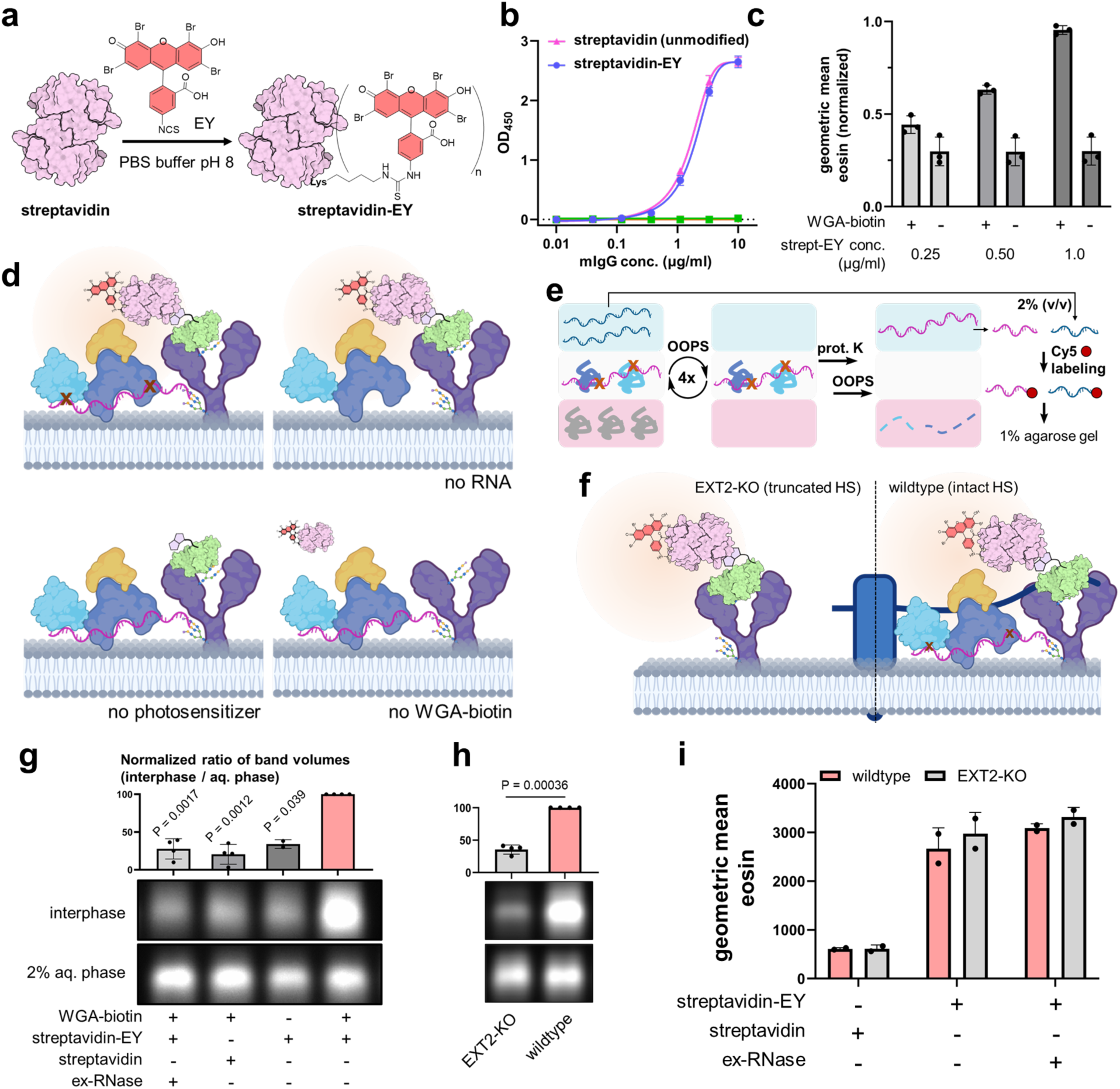
HepRNA HS-dependency validation by TLR7-independent cell-surface crosslinking experiments. **a**) Reaction scheme for eosin isothiocyanate (EY)-streptavidin conjugation. Isothiocyanate generally react with primary amines on proteins, forming thiourea. **b**) Enzyme-linked immunosorbent assay (ELISA) demonstrates modified streptavidin retains full capacity for biotin binding. Blue circle, with streptavidin-EY coated on the plate, incubated biotinylated IgG; pink triangle, unmodified streptavidin, biotinylated IgG; green square, streptavidin-EY, unmodified IgG; orange triangle (inverted), unmodified streptavidin, unmodified IgG. **c**) Titration of streptavidin-EY on live cell surface with and without biotinylated WGA deposition. **d**) Schematics of EY-mediated cell surface crosslinking experiment and controls. In no-RNA control, lives cells were pretreated with RNase I. **e**) Sample processing after live cell surface crosslinking. Abbreviations: OOPS, orthogonal organic phase separation; prot.K, proteinase K; Cy5, for this experiment, cyanine 5-conjugated cytidine-5′-phosphate-3′-(6-aminohexyl)phosphate. **f**) Schematics of EY-mediated cell surface crosslinking on wildtype and HS-deficient cells. **g**) 1% agarose gel electrophoresis of 3’-end Cy5-labeled RNA from samples depicted in panel **d**. Gel image were cropped for small RNA region. Bar graphs show the normalized band volumes corrected by aqueous phase total RNA. n = 4 independent cell cultures; n = 2 for no-WGA-biotin control. Error bars, s.d. **h**) Image and bar graphs of 1% agarose gel electrophoresis of 3’-end Cy5-labeled RNA from wildtype and EXT2-KO (HS-deficient) cells. The wildtype samples were produced in the same batch as EXT2-KO samples. n = 4 independent cell cultures. **i**) Bar graphs of flow cytometry quantification of streptavidin-EY recruited on the surface of wildtype cell with or without RNase treatment and EXT2-KO cells.

The crosslinked csRNA (pink RNA in **Figure 4e**) was then released by proteinase digestion of the AGPC interphase after repetitive washes. The released csRNA was then fluorescently ligated at 3’-end and subjected to agarose gel electrophoresis. A small portion of total RNA (blue RNA in the first aqueous phase in **Figure 4e**) was also taken along and processed in the same fashion to ensure the inputs for AGPC are consistent across different samples. Strong fluorescent signal of released interphase RNA was observed only when both strep-EY and WGA-biotin are present (the fourth lane of **Figure 4g**). Leaving out either component afforded much weaker signals (the second and third lanes). Pretreating live cells briefly with extracellular RNase to degrade csRNA (first lane) resulted in a significantly weakened signal compared to untreated, csRNA-intact cells.

We then employed SCOOPS to validate HS’s essential role in presenting hepRNA-RBP complexes at cell-surface. We expected cells deficient in HS biosynthesis should afford less SCOOPS-isolated RNA due to the absence of the intact polysaccharide chain as a scaffold for hepRNA presentation (**Figure 4f**). Indeed, we observed a significant reduction of SCOOPS-isolated RNA fluorescence from HS-deficient cells as compared to the wildtype in the agarose gel (**Figure 4h**). To ensure the reduced signal was caused by the biology rather than differences in SO generator recruitment on cell surface, we performed flow cytometry assays to quantify recruited strep-EY on wildtype and HS-deficient cells (**Figure 4i**). We found strep-EY introduction to cell surface was at a comparable level between wildtype and HS-deficient cells. Taken together, the results demonstrated that there is a portion of SCOOPS-isolated RNA which becomes largely absent on HS-deficient cells, suggesting hepRNA’s dependency on the intactness of HS for a stable presentation on cell surface. Thus, SCOOPS validated the observation made with the TLR7 probe in an orthogonal manner.

### hepRNA identification by sequencing of SCOOPS-isolated RNAs

Provided that hepRNA could be potentially captured by SCOOPS, we sought to identify these RNA species using next-generation sequencing. Bioanalyzer traces revealed that SCOOPS-isolated RNA in wildtype cells had a wide length distribution (**Supplementary Figure 8a**), with a bulge in electropherograms spanning a large range of recorded length. Such bulge was much less prevalent in the HS-deficient mutant. The electropherograms of SCOOPS-isolated RNA in wildtype cells were vastly different from total RNA, as the characteristic peak pattern as observed in the latter was almost lost. To cover both polyadenylated and non-polyadenylated RNA biotypes in the sequencing library, we took a ligation-based strategy to introduce sequencing adapters (**Supplementary Figure 8b**). The resulted product was reverse-transcribed, ligated with a second adapter, amplified using polymerase chain reaction (PCR). The PCR product then underwent polyacrylamide gel-based size selection (**Supplementary Figure 8c**). The size-selected libraries were subjected to Illumina platforms for sequencing.

The analysis of sequencing data revealed large difference between SCOOPS-isolated RNA and the total transcriptome (see **Supplementary Table 6** for a complete RPM table), as demonstrated by the principal component analysis (**Supplementary Figure 9e**). Comparing SCOOPS-isolated RNA with total RNA (SCOOPS_WT vs. total_WT) can identify the transcripts which were particularly well retained in the interphase due to crosslinking. These transcripts are colored as yellow dots in the scatter plot (**Figure 5a**). It is important to note that not all RNAs in the interphase were crosslinked ones on the cell surface: as demonstrated in the biochemical assays (**Figure 4g**) and bioanalyzer results (**Supplementary Figure 8a**), there were also background RNA species retained in the interphase even without crosslinking (see **Supplementary Text 3** for a detailed explanation).

**Figure 5.**
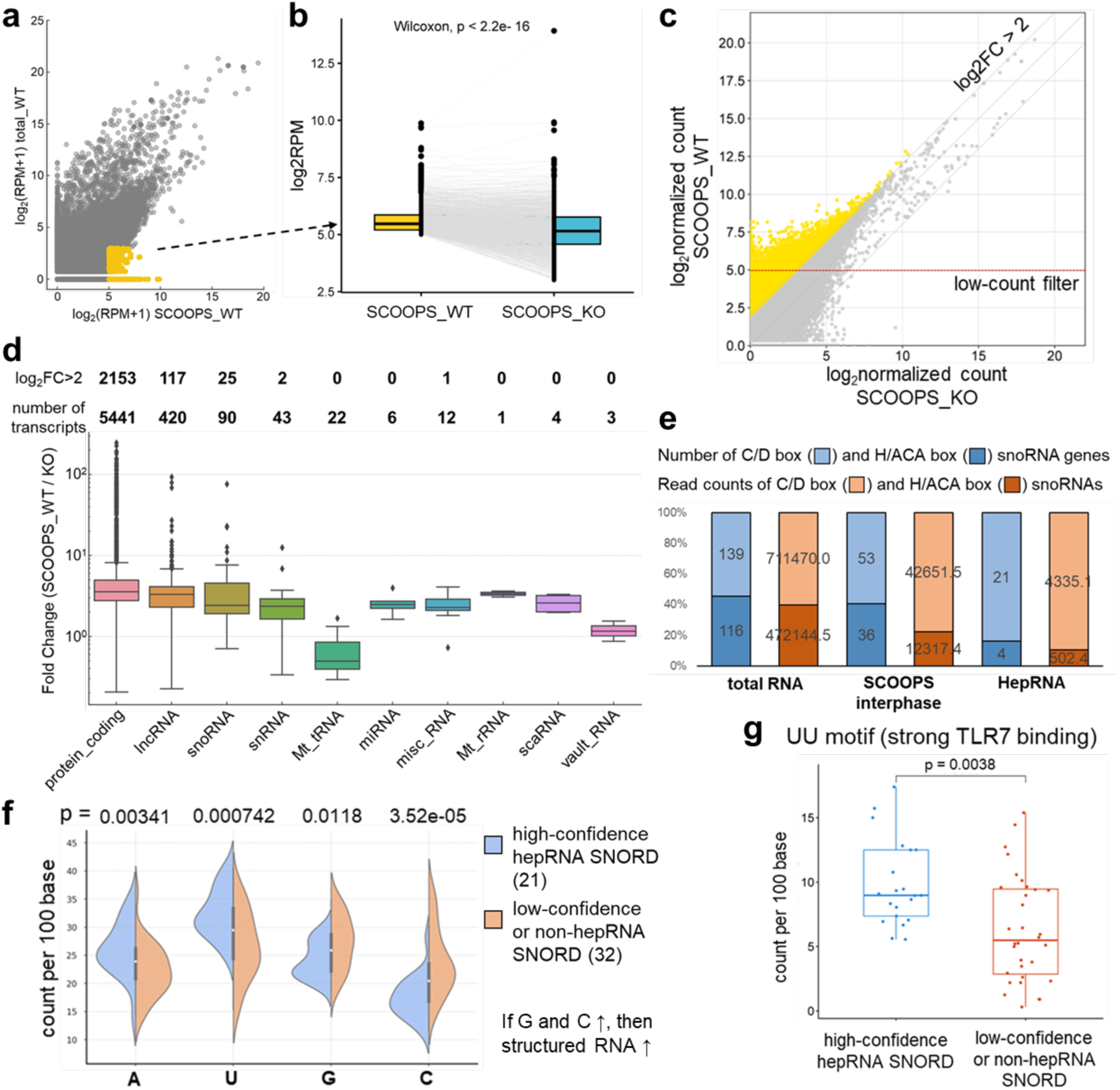
HepRNA identification via next-generation sequencing. **a**) Scatter plot of mean RPM values. n = 2 independent SCOOPS experiments. Mean (RPM+1) was log2-transformed. Transcripts are displayed without low-count filtering. X-axis is SCOOPS_WT sample, and Y-axis is total cellular RNA from wildtype cells. **b**) Connected box plot of RPM of SCOOPS interphase-retained transcripts in wildtype (SCOOPS_WT) and EXT-KO (SCOOPS_KO) samples. Grey lines connect corresponding transcripts in both datasets. Statistical significance, Wilcoxon test. **c**) Scatter plot of mean read counts normalized on background RNA species, comparing SCOOPS_WT (Y-axis) with SCOOPS_KO (X-axis). High-confidence hepRNA (yellow dots) defined as the transcripts which have 4 times higher read counts post-normalization (on background RNA) in SCOOPS_WT than in SCOOPS_KO. Transcripts with normalized read counts larger than 32 in SCOOPS_WT were taken for further analysis in panel **d** – **g**. **d**) Box plot summarizing fold change values per RNA biotype derived from SCOOPS_WT over _KO. **e**) Bar graph illustrating relative composition of snoRNA subtypes in total transcriptome, SCOOPS interphase, and as high-confidence hepRNA. **f**) Violin plot showing significant difference in nucleotide compositions between high-confidence hepRNA and low-confidence/non-hepRNA SNORD transcripts. Mann-Whitney U test was used to derive p values for each nucleotide. **g**) Box plot showing strong TLR7-binding motif (UU) occurs more frequently in high-confidence hepRNA SNORD transcripts than in low-confidence/non-hepRNA ones. Statistical significance, Mann-Whitney U test.

We asked if any hepRNA had been retained in the SCOOPS interphase. We define hepRNA as transcripts that are substantially reduced or absent in the SCOOPS interphase of EXT2-KO mutant (SCOOPS_KO), when compared to those in the wildtype SCOOPS interphase (SCOOPS_WT). As illustrated in the connected box plot (**Figure 5b**), of the well-retained transcripts in the SCOOPS_WT interphase (yellow dots in **Figure 5a**), many had substantially reduced counts in SCOOPS_KO samples. These transcripts include mainly mRNA and lncRNA, which have been major csRNA biotypes in previously reported datasets.^3, 4^ In particular, MALAT1, as one of the few validated cell-surface lncRNA, was also highly enriched in the SCOOPS interphase as hepRNA. The results suggested SCOOPS effectively captures RNA at cell surface, and hepRNA is indeed present in the SCOOPS interphase.

To comprehensively identify hepRNA candidates, we first normalized the read counts of SCOOPS_WT and SCOOPS_KO on the background RNA species (see **Supplementary Text 4** for definitions and rationale). It is crucial to note that while SCOOPS is designed to crosslink all RNA-protein complexes at the site of SO generation, it is not specific to hepRNA. The key to ensuring hepRNA specificity was the use of HS-deficient cells as a negative control. In this setup, RNA species that are 1) present at the cell surface but independent of HS or 2) non-specifically crosslinked or isolated during SCOOPS would show minimal enrichment when comparing SCOOPS_WT with SCOOPS_KO datasets.

A higher fold change (FC > 4, SCOOPS_WT over _KO) indicates that a transcript’s cell-surface presentation is strongly dependent on HS rather than other mechanisms, classifying it as a high-confidence hepRNA candidate. This criterion applies to many mRNA, lncRNA, and snoRNA transcripts (**Figure 5c** and **d**, see **Supplementary Table 7** for a full candidate list). In contrast, transcripts with a lower fold change (2 < FC < 3) are considered low-confidence hepRNA candidates, as they may associate with HS only partially or could partly result from non-specific crosslinking or isolation during SCOOPS. These transcripts span nearly all investigated biotypes. However, mitochondrial tRNA (Mt_tRNA) and vault RNA are unlikely to be hepRNA, as their transcripts were barely enriched in the SCOOPS_WT vs. SCOOPS_KO comparison.

While mRNA and lncRNA biotypes dominate hepRNA candidates, which is consistent with previous studies, strikingly, snoRNAs also constitute a considerable portion of hepRNA. Among the high-confidence snoRNA-derived hepRNA candidates, the C/D box type outnumbers the H/ACA type (21 SNORD vs. 4 SNORA), both in transcript count and total read coverage (**Figure 5e**). Remarkably, we identified defining structural traits that distinguish high-confidence SNORD hepRNAs (hepSNORDs). Firstly, high-confidence hepSNORDs contain significantly less guanosine (G) and cytidine (C) compared to low-confidence/non-hepSNORDs (**Figure 5f**). Such nucleotide composition suggests high-confidence hepSNORDs may harbor less duplex or structured region intra- or intermolecularly.^49, 50^ Furthermore, in high-confidence hepSNORDs, the occurrence of consecutive uridines (UU), a motif known for strong TLR7 binding,^20^ is significantly more frequent than low-confidence/non-hepSNORDs (**Figure 5g**). This compelling observation reveals that specific RNA primary structures underlie the ability of this subset of RNA to associate with cell-surface HS. Beyond structural implications, our findings based on a TLR7-independent method provide an explanation at RNA primary structure level for TLR7’s selectivity for hepRNA.

### hepRNA recruits immune receptor on cell surface

The vicinity of hepRNA to cell surface glycoproteins cell surface motivated us to explore the possibility that ligands or receptors to these proteins may be regulated by hepRNA. Our hepRNA-proximal datasets harbor many membrane glycoproteins which were no longer enriched in response to RNase treatment (**Figure 6a**). In search for an appropriate hit to follow up on from our datasets, we found glycoprotein poliovirus receptor (PVR, or CD155) had a decrease of ∼28 folds in its intensity in RNase dataset compared to that in TLR7 one, suggesting PVR is spatially close to hepRNA. PVR was also a common hit between the hepRNA-proximal and our biotinylation-OOPS datasets (**Supplementary Figure 6g**). Consistently, in a previous study, the mobility of PVR from cell lysates through a sucrose density gradient dramatically changed upon RNase treatment.^51^ Additionally, PVR is documented in RBP2GO database, despite with a low RBP2GO score.^52^

**Figure 6.**
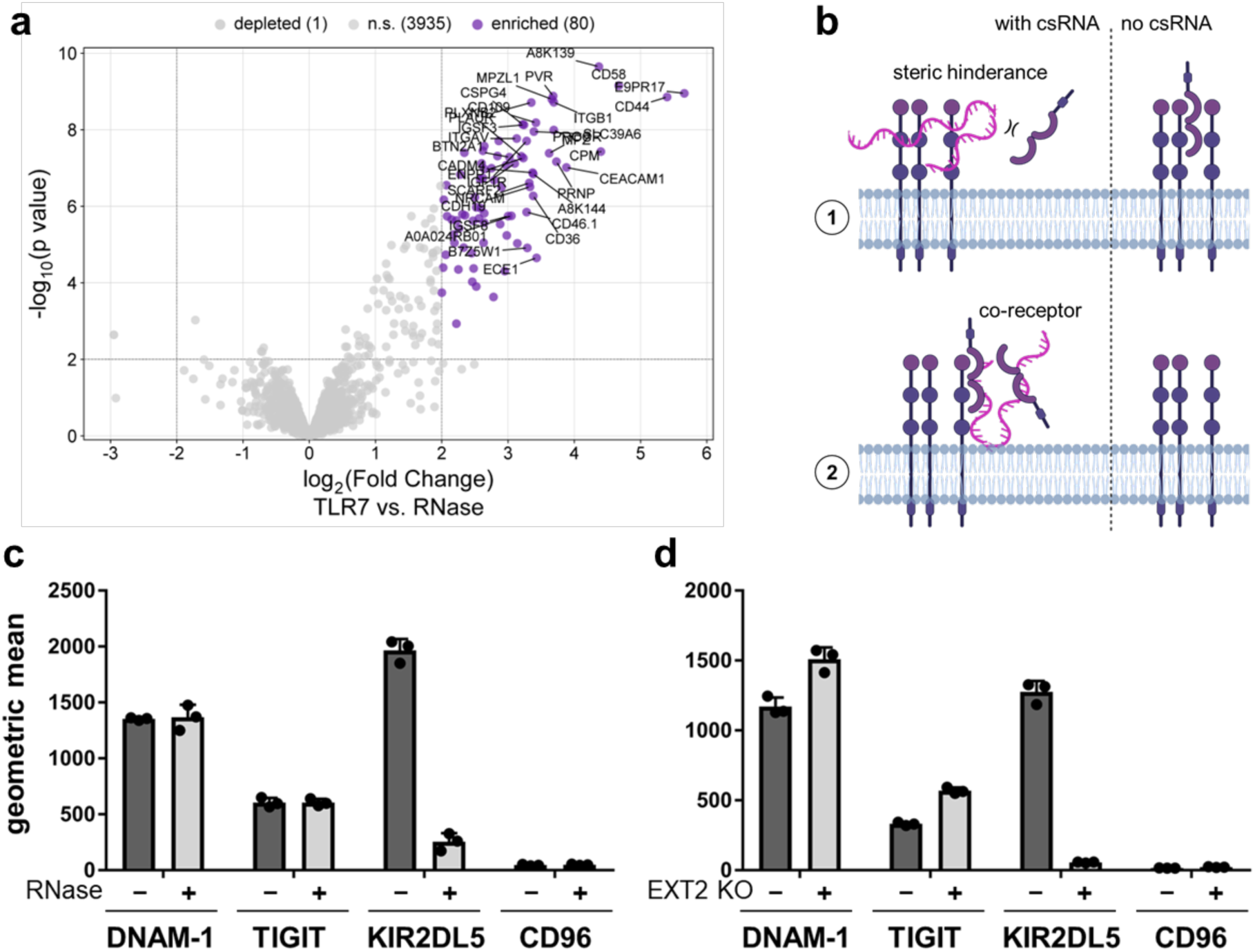
HepRNA recruits KIR2DL5 on cell surface. **a**) Volcano plot from the comparison of proteome between TLR7-mediated proximity labeling with vs. without RNase treatment. Significantly enriched proteins with high fold change (≥ 8, p < 0.01. n = 3 independent cell cultures) are labeled with their names. **b**) Schematic representation of how csRNA may regulate ligand-receptor interactions on the plasma membrane. **c**) Bar graphs from flow cytometry analysis of live cell surface binding of PVR-binding Fc-fused recombinant proteins, in the presence or absence of csRNA. 40 μg/mL RNase A was used to deplete csRNA. AlexaFluor647-conjugated goat-anti-human was used to detect Fc-fused proteins on live cell surface. n = 3 independent cell cultures. **d**) Bar graphs from flow cytometry analysis live cell surface binding of PVR-binding Fc-fused recombinant proteins. Wildtype Mel526 and the EXT-KO mutant cells were compared side-by-side. n = 3 independent cell cultures.

Given the RNA proximity or direct interaction, PVR was thus selected for further investigation. PVR overexpresses in different cancers and can bind to T cell immunoreceptor with immunoglobulin and ITIM domain (TIGIT),^53^ CD96,^54^ and Killer cell immunoglobulin-like receptor 2DL5 (KIR2DL5),^55^ all of which are inhibitory receptors able to suppress the killing by T cells or NK cells. PVR also interacts with an activating receptor, CD226.^56^ The engagement of PVR with these proteins forms immune checkpoints, and they are thus new promising targets for cancer immunotherapy. We have envisioned the possible modes of action of HepRNA on PVR recognition (**Figure 6b**): 1) hepRNA may sterically hinder the binding; 2) hepRNA may facilitate the binding of specific proteins to PVR by functioning as a co-receptor, or by priming PVR for optimal recognition; and finally, 3) hepRNA may be just a bystander, and has no effect on binding.

We used flow cytometry to quantify the RNase responsiveness of cell-surface binding by the soluble, Fc-tagged version of ectodomains from the four known PVR-binders with and without RNase pre-treatment (**Figure 6c**). TIGIT and CD226 exhibited little change in response to RNase treatment, indicating that HepRNA does not modulate their recognition. CD96 did not detectably bind to either cell surface in our experimental settings, suggesting additional factors may be required. KIR2DL5 did show a dramatic reduction in fluorescent signals on the cell surface when the live cells were pretreated with RNase. In line with the genetic screening data, the EXT2 KO mutant cells also exhibited a strong reduction in KIR2DL5 binding, but not for CD226 or TIGIT (**Figure 6d**). Neither the RNase treatment nor EXT2 KO negatively affected cell-surface PVR levels (**Supplementary Figure 10a** and **b**), indicating that the decrease in KIR2DL5 binding was due to hepRNA removal, rather than an effect on PVR expression. Exogenous RNA added to prior hepRNA-depleted cells restored partly KIR2DL5 binding (**Supplementary Figure 10c**), suggesting RNA-KIR2DL5 interaction played an important role in recruiting the latter on cell surface.

To demonstrate KIR2DL5 binds to hepRNA on living cells, we transiently overexpressed the full-length protein fused with FLAG tag on wildtype and HS-deficient mutant (pgsD-677) CHO cells (**Supplementary Figure 10e**. See **Supplementary Figure 4e** for hepRNA detection). We expect the expression of human KIR2DL5 in such a non-human background should minimize protein-protein interactions. Upon KIR2DL5 overexpression, the cells were exposed to UVC irradiation to crosslink RNA with bound proteins in situ. After cell lysis, we performed immunoprecipitation against FLAG tag to pull down KIR2DL5, and if any, the crosslinked RNA bound to the protein. The co-precipitated KIR2DL5-bound RNA was enzymatically ligated with biotin at 3’-end on-bead and finally subjected to Western blot for the detection of biotin. Strong biotin signal as a smear was found above recombinant FLAG-tagged KIR2DL5 band, suggesting RNA co-precipitation (**Supplementary Figure 10e**). Such smear disappeared in HS-deficient CHO cells, which indicated an HS dependency of KIR2DL5-bound RNA. To confirm it was the extracellular domain of KIR2DL5 that mediated RNA binding, we performed in-vitro assays in which human Fc-fused KIR2DL5 extracellular domain or IgG isotype control was incubated with 3’-end biotinylated total cellular RNA fragments, UVC crosslinked, and subjected to Western blot. Strong signal was found only for KIR2DL5 crosslinked with biotinylated RNA, whereas that in IgG control was hardly detectable (**Supplementary Figure 10f**). In addition to experimentally demonstrating hepRNA can interact with KIR2DL5’s extracellular domain, we also postulated a structural model for the interaction using AlphaFold 3^57^ (see **Suplementary Text 5** and **Supplementary Figure 11** for a detailed explanation).

To confirm the binding of KIR2DL5 proteins to the cell surface is at least partly dependent on PVR, we generated PVR KO cells using CRISRP-Cas9. Upon treatment with extracellular RNase, the residual KIR2DL5 binding was significantly lower in PVR KO cells than in the wild type (**Supplementary Figure 10d**). This indicated that KIR2DL5-PVR interaction accounted for a portion of fluorescence signals detected on the cell surface. Taken together, these results demonstrate that hepRNA alone can recruit KIR2DL5 on the cell surface, whereas PVR alone only weakly binds to soluble KIR2DL5. The data supports the model that hepRNA functions as a co-receptor for KIR2DL5 and thereby facilitates its engagement with PVR by increasing local KIR2DL5 concentration at cell surface.

## Discussion

### A hepRNA-csRBP-HS ternary complex model

Taking integrated omic-wide approaches and using multiple orthogonal validatory methods, our study answered the pressing question in the emerging field of RNA cell-surface localization: how the biomacromolecule is associated with the plasma membrane. The study was commenced by an exploitation of nature’s own RNA sensor, TLR7, as a probe to detect RNA at the cell surface. The use thereof in a genome-wide CRISPR-Cas9 knockout screening led to the discovery of HS polysaccharide as an essential molecular factor for RNA’s association with the cell surface. While direct interactions between carbohydrates and nucleic acids have been reported,^58^ our data supports the presence of a number of RBPs at cell surface as crucial adapter molecules to bridge the two highly negatively charged biopolymers, forming hepRNA-csRBP-HS ternary complexes (**Figure 7**).

**Figure 7.**
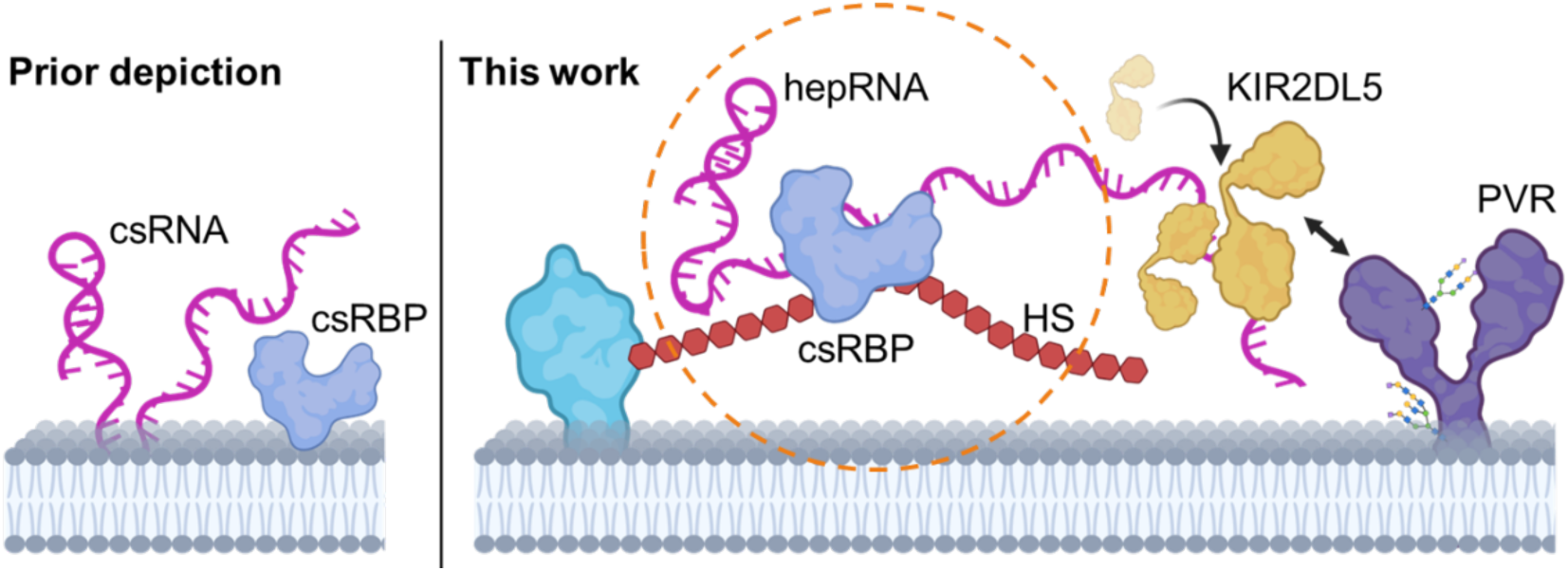
Schematic representation of advanced understanding cell-surface RNA biology. The hepRNA-csRBP-HS ternary complex model is shown in dashed orange circle.

In addition to the routine validatory experiments, such as immunofluorescence, flow cytometry, and Western blot assays, we have established multiple orthogonal methods to validate such ternary molecular configuration. We considered that the csRBPs had been an essential element collectively for RNA’s association with HS, but functionally redundant as individual proteins, particularly as none of the csRBPs from proteomics study were hits in the genetic screening. This was because the knockout of one RBP during the pooled screening may have been compensated by others, such that little, if any, apparent decrease of global hepRNA could be observed in flow cytometry using the TLR7 probe. Due to this collectively essential but individually redundant nature of csRBPs for hepRNA, knocking out these RBPs altogether until a global reduction in hepRNA is seen would fall experimentally impractical. We, therefore, broke down the validation of the termolecular model into separate steps.

Firstly, we sought to validate, in a high-throughput manner, the involvement of hepRNA-proximal csRBP candidates in RNA-RBP complex formation on plasma membrane. This was done by a TLR7-independent cell-surface biotinylation method, combined with UVC crosslinking and the isolation of RNA-bound proteins using OOPS. While 41 hepRNA-proximal proteins were validated as RNA-bound csRBPs, the other hits uniquely identified in biotinylation-crosslinking-OOPS experiment suggested there are more csRBPs than hepRNA-bound ones, because the biotinylation chemistry and UVC crosslinking are not hepRNA-specific. These proteins may be presented on cell surface via an HS-independent mechanism, such as protein-protein interactions.^9, 59^ A more comprehensive mechanistic understanding of csRBPs will likely unveil new biology at cell surface, and warrants future studies.

We then introduced SCOOPS, a photocatalytic spatio-selective crosslinking technology to selectively capture cell-surface RNA-protein complexes. SCOOPS is independent of any RNA-binding probes such as TLR7. Thus, the essential role of HS as a scaffold for hepRNA-csRBP complexes could be validated using an orthogonal approach. Furthermore, SCOOPS enabled unveiling the identities of hepRNA for the first time. It revealed that decreased G and C content and a more frequent UU motif occurrence are traits in the RNA primary structure related to HS association. Apart from sequence motifs, more factors such as their secondary and higher-order structures and the transcript-specific interacting proteome should be examined, and how they function in an intertwined manner to regulate RNA cell-surface localization remains to be elucidated.^60^

While our data supports a ternary complex model of hepRNA-csRBP-HS, the structural basis for the interaction between RBPs and HS remain elusive. Thus, future work will be needed on investigating the molecular details how RBPs bind to HS. HS is known to be a molecular mimicry of nucleic acids and there have been reports showing several nucleic acid-binding proteins can indeed interact with HS. Generally, the interaction between RBP and HS may be mediated by a designated HS-binding domain present on the RBP, or a promiscuous RNA-binding domain,^61^ which typically harbors positively charged side chains for the electrostatic interaction with HS. SSB, for example, contains a putative HS-binding motif (GRRFKG) at its C-terminus disordered region, implying potential HS-binding activity.^62^ Conversely, HNRNPA1 does not contain any known HS-binding motif, despite its presence on the cell surface of wildtype, but not on HS-deficient mutant cells. It suggests a yet-to-be-characterized mode of interaction between HS and the RNA-binding domains or other portion of the protein.

### hepRNA as a co-binder for KIR2DL5

We further demonstrate that once displayed on the cell surface, the RNA can enrich glycoprotein immune receptors, likely to facilitate the interaction with their cognate ligands. In the case of KIR2DL5, hepRNA functions as a co-binder, such that local concentration of KIR2DL5 becomes much higher, thereby facilitating PVR engagement, and compensating for the weak affinity of PVR for KIR2DL5.^55^ The co-binder effect in modulating cell surface receptor-ligand interactions is not uncommon. For example, the engagement of fibroblast growth factor 2 (FGF-2) with FGFR requires the former to engage with HS, likely resulting in a liquid-liquid phase separation.^63^

KIR2DL5, together with a few other KIR family members, has been previously shown to bind to heparin in-vitro via the d0 domain, which is rich in positively charged amino acids.^64^ Heparin is often considered a molecular mimicry of nucleic acids because of its heavy negative charges along the polysaccharide chain and has been used to purify select nucleic acid-binding proteins. Although binding to heparin in-vitro, we demonstrated that KIR2DL5 could directly bind to RNA in-vitro and, at least partly, on living cell surface, for which a structural basis was predicted using AF3. Notably, such promiscuity in binding has been described for a cell-surface-localized protein, receptor for advanced glycation endproducts (RAGE). It was shown previously to bind to HS,^65^ and more recently also to DNA.^66^ Similar observation has also been made for certain antibodies directed against DNA, which can cross-react with HS.^67, 68^

### Origin of hepRNA

One key question remains as to the origin of the hepRNA and the associated csRBPs. It is plausible that the RNPs are transported from cytosol to the plasma membrane of the same cell (“inside-out”). However, the “inside-out” model requires the translocation of cytosolic RNA and RBPs to the luminal side of specific membraneous organelles.

Interestingly, a recent study^69^ showed a distinct set of RNAs from the cytosolically localized transcripts can be found in the ER lumen, implying an unidentified pathway for RNA luminal translocation. In our genetic screening study, TMED10 was identified as an essential factor for hepRNA presentation. This multifunctional protein forms channels on the ERGIC membrane and mediates the translocation of select cytosolic proteins without leader peptide to the luminal side for secretion.^70^ Of particular interest, TMED10 was reportedly not relevant to HS biosynthesis^71^ and has been documented to potentially bind to RNA in the RBP2GO database,^52^ implicating their possible roles in vesicular RNA transport.

An equally likely scenario would be that they are secreted from live cells or released from dead/dying cells and directly deposited on the HS chains (“outside-on”). There are numerous biological processes in which intracellular components including RNA (protein unbound or in complex) are released to extracellular space. The release can be from dying/dead cells or damaged tissues, but also living cells.^72^ Extracellular RNAs or their protein complexes are common damage-associated molecular patterns (DAMPs) which can activate innate and adaptive immunity and are associated with many disorders,^73^ such as sepsis,^74^ cancer,^75^ and autoimmune diseases.^76^ The observation that HS, but not other proteoglycans, can selectively capture extracellular RNA-protein complexes is strong evidence of a specific and biologically relevant interaction, suggesting that HS plays a unique role in regulating extracellular RNA dynamics and immune activation: On the one hand, it may limit RNA-based DAMPs locally by capturing them to slow down their lateral diffusion; on the other hand, it may also serve as a scaffold to present the RNPs to immune cells for activation. Further exploration of HS-RBP-RNA interactions could uncover novel regulators of immune signalling in tissue injury, offering new insights into inflammation-related diseases.

## Supplementary Text 1. Additional insights from genome-wide screening

### RBPs as hits in TLR7^low^ subpopulation

Among the hits from the genome-wide screening, there are a few putative RBPs (see **Supplementary Table 2** for a complete list). ATP-dependent RNA helicase **DHX29** (RBP2GO score 43.1) is a poorly understood classical RBP involved in translation initiation.^77^ Transmembrane emp24 domain-containing protein 10 (**TMED10**) is a multifunctional protein in the cytoplasm. It functions as a cargo receptor involved in protein vesicular trafficking and quality control in ER and Golgi. It can also oligomerize and act as a protein channel to facilitate the post-translational entry of leaderless cytoplasmic cargo into the ER-Golgi intermediate compartment (ERGIC) for extracellular secretion.^70^ TMED10 is also a non-canonical RNA-binding protein with an RBP2GO score of 9.7. The function of the hit appears interesting, as the ribonucleoproteins do need to be transported from cytosolic side to luminal side of organelles or vesicles to become topologically connected to plasma membrane. UDP-glucose 6-dehydrogenase (**UGDH**) is an enzyme which creates sugar building blocks for proteoglycan synthesis, and is a non-canonical RNA-binding protein (RBP2GO score 3.8). The RNA-binding activity is expected to be unrelated to csRNA biology. Transmembrane 9 superfamily member 3 (**TM9SF3**) is a poorly understood protein regulating glycosyltransferase function in Golgi, and is a non-canonical RNA-binding protein (RBP2GO score 5.7).

Interestingly, CD44, a glycoprotein known to be carrier protein of HS polysaccharide,^78^ was a hit (criteria, p < 0.01 AND at least two guide RNA enriched) in one biological replicate of the genome-wide screening. In this replicate, it had an enrichment score of 4.40, with 2 out of 4 guide RNAs significantly enriched. In the other replicate, CD44 had one guide RNA significantly enriched, the gene had a moderate enrichment score of 1.27. Moreover, CD44 has been documented in the R-DeeP database^51^ to be potentially associated with RNA, either directly as an RBP or indirectly as a complex with other RBPs.

### Hits from TLR7^high^ subpopulation

After the two-round phenotypic selection using fluorescence-assisted cell sorting (FACS), we have observed a clear bimodal distribution for TLR7-bound subpopulation, with a small but distinct TLR7-negative subpopulation (**Supplementary Figure 3**). However, there was no such distinct subpopulation with particularly high TLR7 signal even after the second phenotypic enrichment. The significantly enriched hits consistent in two biological replicates included N-acylneuraminate cytidylyltransferase (**CMAS**), Uroporphyrinogen decarboxylase (**UROD**) and C-type lectin domain family 17, member A (**CLEC17A**). Despite CMAS and UROD are documented in the RBP2GO database, their relevance to csRNA remain clear.

## Supplementary Text 2. Observation of singlet oxygen-mediated RNA-protein crosslinking

The small molecule photosensitizer, eosin Y (EY), is known to passively diffuse into living cells, and has been used to generate singlet oxygen (SO) in cells with the irradiation with green light (**Supplementary Figure 7a**).^48^ When performing acid guanidinium thiocyanate-phenol-chloroform (AGPC) extraction of total RNA (from the blue 1^st^ aqueous phase in figure panel **a**, without repeated OOPS), we observed substantial thickening of the interphase, and a dropped yield of RNA isolated from the aqueous phase (**Supplementary Figure 7b**). We reasoned the RNA must have been brought to the interphase (off-white phase in panel **a**), a similar effect which occurs when the cells are exposed to UVC irradiation.^42^ UVC irradiation creates covalent bonds between the RNA and the bound proteins, making the complexes into an amphiphilic entity. This physical nature of crosslinked RNA-protein complexes retains them in the AGPC interphase. Based on our observation, EY-generated SO may have led to RNA-protein crosslinking, creating the amphiphilic species.

We performed OOPS on EY-treated, green light irradiated cells (denoted EY in figures), in parallel with UVC irradiated cells (UV) and a no crosslinking control (NC). The OOPS-cleaned interphase can be treated with proteinase K to degrade bound proteins, releasing the crosslinked RNA back to the aqueous phase. The released protein-bound RNA was isolated from aqueous phase by another round of AGPC. Agarose gel electrophoresis (SybrGold stain) showed that there was indeed RNA in the interphase of UV and EY samples, with a corresponding reduction of the amount of RNA in the first aqueous phase (**Supplementary Figure 7c**). Most total cellular RNA could be isolated from the first aqueous phase of NC samples.

If RNA were crosslinked to bound proteins by EY-generated SO, we should also expect the presence of RNA-binding proteins (RBP) in the interphase (**Supplementary Figure 7d**). Treating the OOPS-cleaned interphase with an RNase cocktail releases RNA-bound proteins to organic phase (pink phase). We performed SDS-PAGE for the released RNA-bound proteins from EY and UV samples, followed by a Coomassie gel stain (**Supplementary Figure 7e**). Stained bands from EY samples were substantially stronger than those of UV, suggesting more crosslinking events may have occurred for EY than UV. We also performed Western blot to detect known RBPs, including heterogeneous nuclear ribonucleoprotein A1 (HNRNPA1), enolase 1 (ENO1), Sjögren syndrome antigen B (SSB) (**Supplementary Figure 7f**). HNRNPA1 gave strong bands in EY samples, and moderate signals in UV. ENO1 and SSB did afford strong to moderate signals in EY, but only weak to undetectable signals in UV. The results suggest EY can crosslink RNA to proteins and the amphiphilic entities can be brought to the AGPC interphase for isolation. Take together, these results serve as technical basis for the development of SCOOPS.

## Supplementary Text 3. Detailed discussion on the bioanalyzer electropherograms

The electropherograms of SCOOPS_KO samples indicate the SCOOPS interphase contains more RNA components than exclusively hepRNA. Two prevalent peaks observed in the bioanalyzer traces of SCOOPS_KO samples include one in small RNA region (fewer than 200 nucleotides), and the other aligned well with 28S ribosomal RNA (rRNA) (**Supplementary Figure 8a**). The peaks may be given rise to by RNA species exhibiting an intrinsic physical property to be retained in the interphase regardless of whether SO-crosslinked. The SCOOPS interphase could also contain originally intracellular RNA from, for instance, cell debris, which could be randomly crosslinked by singlet oxygen generated at cell surface. In addition, the interphase may also contain cell-surface localized RNA species whose presentation is not dependent on HS, which is out of the scope of this study. Importantly, SCOOPS_KO samples can provide indications for hepRNA-irrelevant RNA species intrinsically associated with the SCOOPS procedure. These RNA species can serve as background in RNA sequencing data analysis (discussed in detail in **Supplementary Text 4**).

The bioanalyzer traces of SCOOPS_WT samples revealed that SCOOPS interphase contains RNA species with a wide length distribution. The content of the interphase became much less when intact heparan sulfate (HS) chain is missing. This is in consistency with the biochemical assays using in-gel fluorescence. While the bioanalyzer traces gave a more global view of the length profile of SCOOPS-isolated RNA, the major difference in RNA in-gel fluorescence (**Figure 4h**) between SCOOPS_WT and _KO was observed only for small RNA region. This is likely due to substantially higher numbers of exposed 3’-end in small RNAs for fluorophore ligation catalyzed by T4 RNA ligase 1, while long transcripts have much fewer ligation sites available for the enzyme.

## Supplementary Text 4. Read count normalization on background RNA species

### Rationales

As indicated by in-gel fluorescence (**Figure 4h**) and Bioanalyzer electropherograms (**Supplementary Figure 8a**), SCOOPS_KO contains background RNA species intrinsically associated with the SCOOPS procedure, whereas SCOOPS_WT contains hepRNA as well as the background RNA. Thus, the background RNA could serve as an internal standard and provide a means to normalize sequencing data. This is similar to the idea of normalization on a minor set of genes/transcripts that remain unchanged between biological conditions, such as housekeeping genes and spike-in controls.^79^ Unlike the routinely used median of ratios^80^ or trimmed mean of M values^81^ methods, which presumes only a small fraction of genes/transcripts differs between conditions, normalization on minor unchanging components is performed when a global shift is expected, which is the case for SCOOPS_WT and SCOOPS_KO. If SCOOPS were performed with comparable cell numbers for WT and KO, the background RNA in SCOOPS interphase, which is intrinsic to the technique, should be of similar quantity in the two samples. Due to the presence of hepRNA, WT had substantially more RNA in the SCOOPS interphase (Stage 1, **Supplementary Figure 9a**). However, equitizing masses of RNA input for sequencing library preparation led to the reduced background RNA present in SCOOPS_WT than in SCOOPS_KO (Stage 2). Such difference in background RNA species in WT and KO were kept throughout the entire process, particularly as both samples underwent the same number of PCR cycles (16 cycles), and were pooled in equimolar for sequencing (Stage 3). Upon normalization by library size (read count per million mapped reads, or RPM), the background RNA species in SCOOPS_WT should consistently have smaller RPMs than those in SCOOPS_KO (Stage 4). To make it a fair comparison between SCOOPS_WT and _KO, the background RNA in the former should be scaled to the same level as in the latter. The scaling factors for normalizing on background RNA should then be applied to all mapped genes/transcripts in each sample (Stage 5).

### Scaling factors for normalization

Although external scaling factors could have been derived from comparing the amount of sequencing library input (150 ng) as fraction out of the total amount of SCOOPS isolated RNA in WT and KO samples, we deliberately avoided normalization by these external factors. Instead, we sought to derive the scaling factors internally, by comparing background RNA transcripts between SCOOPS_WT and KO datasets. We define background RNA as those with more than 2 times decrease in their mean RPMs in SCOOPS_WT compared to the _KO sample (log_2_FoldChange[WT/KO] < −1, with a log_2_RPM ≥ 5 filter. See *Rationales*). Of this subset of transcripts, Pearson correlation analysis revealed high similarity (R = 0.97) between SCOOPS_WT and _KO (**Supplementary Figure 9b**), which fits with the hypothesis that the background RNA is intrinsically associated with the SCOOPS procedure and remains unchanged between biological conditions. Library-specific scaling factors were then derived from performing median of ratios normalization for each SCOOPS sample (**Supplementary Figure 9c**). Remarkably, such internal scaling factors correlate well with the external factors (**Supplementary Figure 9d**). The RPMs for all transcripts in each individual library were then uniformly corrected by the corresponding internal scaling factor.

## Supplementary Text 5. AlphaFold3-predicted model for KIR2DL5-RNA interaction

### Structural basis of KIR2DL5-RNA interaction

To provide a plausible structural basis for KIR2DL5-RNA interaction, we employed Alpha Fold 3 (AF3) in generating a structural model for the bimolecular complex. We used stretches of eight oligonucleotide repeats and the extracellular region of KIR2DL5 as the input. The prediction demonstrated that KIR2DL5 ectodomain folded into two distinct domains (namely, d0 and d2) at overall high local confidence (plDDT score) (**Supplementary Figure 11a**), which is consistent with previous reports.^64^ Because the RNA sequences KIR2DL5 binds to have been elusive, we used stretches of repetitive oligonucleotide repeats as RNA input for model prediction. AF3 predicted a moderate to high confidence interaction between KIR2DL5 and RNA (ipTM score) with a well-defined spatial arrangement (low PAE), suggesting that the binding interface is reliable. (**Supplementary Figure 11b** and **c**). Octaguanosine (G_8_) gave 0.88 (between 0 and 1) as the highest ipTM score, suggesting guanosine residues may be important for KIR2DL5-RNA interaction. The predicted model complex shows RNA oligos likely make contacts with the N-terminus d0 domain, but not the C-terminus d2 domain. Consistently, d0 domain harbors a patch of positively charged residues such as lysine and arginine (**Supplementary Figure 11d**), which features are often associated with RNA binding, but further studies would be needed to determine their precise role in this interaction.

### KIR2DL5-RNA interaction specificity

To investigate if KIR2DL5 binding to cell surface may be mediated by any of the cell-surface RBP hits, we also employed AF3 to predict if complexation is likely between KIR2DL5 extracellular domain and several cell-surface RBP hits (see Materials & Methods). Among all tested cell-surface RBPs, none gave an acceptable ipTM score (all below 0.6). Furthermore, we also observed a decrease of local plDDT scores for KIR2DL5 at the predicted interface compared to KIR2DL5 alone. Such worsened plDDT scores and low ipTM values indicate KIR2DL5 is unlikely to interact, at least, any of the screened cell-surface RBPs. Taken together, the computational data, along with the biochemical assay, supports KIR2DL5 directly binds to RNA.

### RNA’s role in regulating oligomerization

We further employed AF3 to gain insights into the molecular function of KIR2DL5-RNA association. KIR2DL5 belongs to the immunoglobulin superfamily (IGSF), and numerous members within this superfamily undergo homophilic interactions.^82, 83^ We asked if homophilic interactions between KIR2DL5 monomers may occur and if RNA might regulate the oligomer/monomer states of KIR2DL5. When KIR2DL5 ectodomains alone as two separate protein entities were input in AF3, the predicted model had an ipTM score of 0.14 (**Supplementary Figure 11e**). We observed substantially reduced local plDDT values at the dimer interface. The high distance errors in the portion off the PAE matrix diagonal line (circled in purple, **Supplementary Figure 11f**) indicate low confidence in the predicted relative positions of the residues across the two separate KIR2DL5 monomers. These results suggest low likelihood for KIR2DL5 ectodomain *alone* to dimerize. However, a stretch of guanosine repeats with varying lengths (G_8_, G_12_, G_16_ and G_20_) as another separate entity in addition to the two protein entities greatly boosted the ipTM score to 0.77 (G_16_) (**Supplementary Figure 11g**). The RNA oligos were predicted to retain contacts with d0 domain of both KIR2DL5 monomers. The local plDDT scores were larger than 90 at the protein dimer interphase, suggesting high-confidence prediction. In stark contrast to protein dimer alone, the off-diagonal portion in the PAE matrix (circled in purple, **Supplementary Figure 11h**) showed low errors, indicating AF3 had higher confidence in predicting the relative positions of the residues across the two separate KIR2DL5 monomers within this ternary complex. Based on the AF3 prediction, we postulate that RNA may facilitate the dimerization of KIR2DL5.

## Materials and Methods

### Cell culture

Mel526, U2OS and HeLa cells were cultured in DMEM (Thermo Fisher Scientific, 31966047) supplemented with 10% Fetal Bovine Serum (FBS, HyClone, SH30071.03HI) and 1x Penicillin-Streptomycin-Glutamine (Thermo Fisher Scientific, 10378016, 100x) at 37℃ with 5 % CO_2_. Wild-type Chinese Hamster Ovarian (CHO) (CHO-K1, ATCC, CCL-61) and heparan sulfate deficient mutant cell lines - CHO pgsE-606 and pgsD-677 (ATCC, CRl-2242 and −2244, respectively) were cultured in Ham’s F-12K medium (Thermo Fisher Scientific, 21127022) supplemented with 10% FBS and 1x Penicillin-Streptomycin and Glutamine at 37℃ with 5 % CO_2_. To minimize possible influence from components in bovine serum, Mel526 cells and the mutant cells were also adapted into 1x Penicillin-Streptomycin-Glutamine-supplemented DMEM containing 1% FBS and 9% chemically defined serum replacement (Panexin CD, Pan-Biotech, P04-93100), following the protocol suggested by the supplier. Prior to experiments, cell culture flasks or dishes were replenished with DMEM containing 10% Panexin-CD without FBS.

### Immunocytochemistry

Cells were fixed 24 hours post-seeding in µ-Slide 8 Well Glass Bottom ibidi chambers (ibidi, 80827,

∼30,000 cells/well) with 4% Paraformaldehyde (PFA) aqueous solution (Electron Microscopy Sciences, 15710-S) in PBS for 10 min at RT and washed with PBS twice. Blocking was performed with blocking buffer containing 3% Bovine Serum Albumin in PBS (Sigma, A7638) for 1 hour at RT in RNase-free conditions by adding SUPERaseIn™ RNase Inhibitor (Thermo Fisher Scientific, AM2696, 1:200). For RNA competition, a synthetic RNA oligo (Dharmacon™) with a sequence 5’-pGUCUUCAAAACUAGGUCGUUUUAGA-3’/biotin/ was added at 10 μM during TLR7-Fc incubation. For RNase-treated samples, RNase Cocktail Enzyme Mix (Thermo Fisher Scientific, AM2296, 1:100) was added to the blocking buffer. Post-blocking, recombinant proteins or primary antibodies were added to cells prepared in the same respective blocking buffer and incubated at 21℃ for 1 hour. Cells were washed 3 times with PBS (2 minutes per wash) followed by the addition of secondary antibodies prepared in the blocking buffer along with DAPI (Roche, 10236276001, 1 μg/mL final concentration) and incubated at 21℃ for 1h. The cells were washed with PBS and imaged with confocal microscopy (Olympus A1R SiM, oil immersion, filter settings for Red (Alexa-546 and others: excitation 551nm, emission 565 nm); Blue (DAPI: excitation 358 nm, emission 463 nm); Green (Alexa-488 and others: excitation 490nm, emission 544nm) and Far-red (Alexa-647 and others: excitation 650 nm, emission 671 nm).

### Electron microscopy

U-2 OS cells were cultured in a 6 cm petri dish at 90-100% confluency. To avoid fixation-induced artefacts, live cells were first stained for csRNA by incubating on ice for 30minutes and then applying TLR7-Fc (3% Bovine Serum Albumin in PBS, final conc 2.5 μg/mL) for 1 hour on ice followed by thorough washing and incubation with PAG-10 (Protein-A-gold 10nm, PAG10 nm/S, OD50, Cell Microscopy Core, UMC Utrecht) solution in PBS (1:50) on ice for 30minutes. Immunogold-labelled cell monolayers were fixed in 1.5% glutaraldehyde in 0.1 M sodium cacodylate buffer for 2 hours before being successively incubated in 1% osmium tetroxide in 0.1 M cacodylate buffer for 1 hour and in 1% uranyl acetate in water for 1 hour. The cells were then dehydrated through a series of incubations in ethanol (70-100%) for 90 minutes and embedded in Epon. The flat embedded cells were sectioned with an ultramicrotome (UC6, Leica, Vienna) using a 35 degrees diamond knife (Diatome, Biel, Switzerland) at a nominal section thickness of 70 nm. The sections were transferred to a formvar and carbon coated 200 mesh copper grid and stained for 20 minutes with 7% uranyl acetate in water and for 10 minutes with lead citrate. EM images were recorded using a Tecnai 12 electron microscope (Thermo Fisher Scientific) equipped with an EAGLE 4k×4k digital camera. For navigation on EM images, montages of images at 11,000× were generated using stitching software.^84^ The stitched images were imaged and annotated using Aperio ImageScope (Leica Biosystems).

### Flow cytometry analysis of csRNA on live cells

Cells cultured to 80-90% confluency were washed once with 1x DPBS after removing culture media, and then lifted with 10 mM EDTA (diluted from 0.5M sterile filtered stock solution) in 1x DBPS at 37 ℃ for 5 minutes. Lifted cells were dispensed into a 96-well FACS plate with 2-2.5 x10^5^ cells per well. For extracellular RNase treatment, PureLink RNase A (Thermo Fisher Scientific, 12091021) were added to the lifting buffer at a final concentration of 40 μg/mL. After spinning down at 300 g for 5 minutes at 4℃ and supernatant removal, cells in each well were resuspended in 30 μL of TLR7-Fc or other Fc-tagged proteins (final concentration 5 μg/mL) in 0.5% BSA in 1x DPBS containing 200U/mL RNasein (Promega, N2511), and incubated on ice for 45 minutes. For secondary only control, cells were incubated with 30 μL 0.5% BSA in 1x DPBS. After incubation, 200 μL cold 1x DBPS were added to each well and the plate was spun down at 300 g for 5 minutes at 4℃. After supernatant removal, cells were resuspended in 30 μL of goat-anti-human IgG AlexaFluor647 conjugate (Thermo Fisher Scientific, A-21445) in 0.5% BSA in 1x DPBS, and incubated in dark for 30 minutes on ice. Upon completion, 200 μL cold 1x DBPS were added to each well and the plate was spun down at 300 g for 5 minutes at 4℃. Finally, cells were resuspended in 200 μL cold 0.5% BSA in 1x DPBS containing 0.1 μg/mL DAPI. The cells were then analyzed on a flow cytometer using HTS settings. Typically, 150 μL cell suspension per well was infused in the system for analysis, with a flow rate of 1 mL/minute.

### TLR7-Fc binding rescue by exogenous RNA

Cells were processed as was described above, except for that after detachment with EDTA buffer containing Purelink RNase A, cells were spun down, resuspended in 1x DPBS containing 200 U/mL RNasein, with varying concentration of total cellular RNA (2 – 0.2 mg/mL), incubated at 4 ℃ for 30 minutes, and washed with cold 1x DPBS, before incubating with TLR7-Fc. The rest of the steps followed exactly the above flow cytometry experiment.

### TLR7-RNA crosslinking, precipitation, and on-bead RNA labelling

The procedure after TLR7-Fc binding follows that of eCLIP with modifications.^29^ Cells in a 15 cm dish at 90% confluency were washed once with 1x DPBS following culture media removal, and was then incubated with 3 mL per dish 10 mM EDTA in 1x DPBS at 37℃ for 5 minutes. Another 7 mL 1x DPBS was added to the dish to wash cells off the dish. The total of 10 mL cell suspension was transferred to a 15 mL Falcon tube. The dish was washed further with 5 mL 1x DPBS, which was then combined in the same tube. The cells were spun down at 300 g for 5 minutes at 4℃, and the supernatant was removed. The cell pellets were taken up in 250 μL TLR7-Fc (final concentration 5 μg/mL) in 0.5% BSA in 1x DPBS containing 200U/mL RNasein, and the suspension was put on ice for 30 minutes, with gentle tapping every 10 minutes. The tube was topped up to 15 mL with cold 1x DPBS, and spun down at 300 g for 5 minutes at 4℃. The pellets were washed twice with 10 mL cold 1x DPBS. The cells in the second wash were directly transferred to a 10 cm dish for UVC crosslinking on ice without the lid using Stratalinker (400 mJ setting). After UVC exposure, cells were transferred back to the tube and spun down. For TLR7 non-crosslinked control, cells after lifting were transferred into a 15 mL Falcon tube and spun down, resuspended in 10 mL 1x DPBS, and transferred to a 10 cm dish for crosslinking. After crosslinking, cells were transferred back to the 15 mL tube and spun down, and incubated with the TLR7-Fc solution, which was followed by two washes.

The pellets from above were lysed in 400 μL lysis buffer (50 mM Tris-HCl pH 7.4, 100 mM NaCl, 1% v/v Igepal CA-630, 0.1% v/v SDS, 0.5% w/v sodium deoxycholate) containing 1x cOmplete protease inhibitor (Roche, 11836153001) cocktail and 1mM PMSF. The solution was sonicated at 20% power for 5 minutes (10s on, 10s off) on ice. To the sonicated lysate was added 2 μL TurboDNase (Thermo Fisher Scientific, AM2238), and 4 μL diluted RNase I (Thermo Fisher Scientific, AM2294, prediluted 1:20 in 1x DPBS). The lysate was then incubated at 37℃ for 5 minutes, followed by the addition of 4 μL SUPERaseIn. To prepare magnetic beads for TLR-Fc precipitation, 10 μL per dish protein A Dynabeads (Thermo Fisher Scientific, 10001D) was washed twice with lysis buffer before added to the above RNA-fragmented lysate. The bead suspension was incubated on an end-to-end rotator at 4 ℃ for 2 hours. The beads were then wash twice with 200 μL CLIP high salt buffer (50 mM Tris-HCl pH 7.4, 1 M NaCl, 1% Igepal CA-630, 1 mM EDTA, 0.1% SDS, 0.5% sodium deoxycholate), three times with 200 μL CLIP low salt buffer (20 mM Tris-HCl pH 7.4, 10 mM MgCl_2_, 0.2% Tween-20, 5 mM NaCl). The precipitated RNA-TLR7-Fc was 3’-end repaired on-bead by incubating with FastAP (Thermo Fisher Scientific, EF0654, 1 U), T4 PNK (NEB, M0201, 20 U), TurboDNase (0.4 U) and SUPERaseIn (5 U) in 10 μL 1xPNK buffer without DTT at 37 ℃ for 30 minutes, followed by the same wash procedure. The end-repaired RNA was subjected to on-bead 3’-end biotinylation by incubating with 20 nM pCp-biotin, T4 RNA ligase high-concentration (NEB, M0437M, 30 U) and SUPERaseIn (5 U) in 1x RNA ligase buffer without DTT, containing 1 mM ATP, 3.6% DMSO, 0.025% Tween-20 and 18% w/v PEG-8000 16 ℃ overnight, which was followed the same wash procedure. After the final wash, beads were directly taken up in 1x LDS sample loading buffer containing 1x sample reducing solution, denatured at 75 ℃ for 10 minutes. The samples were resolved on an NuPAGE™ 4 −12% bis-Tris mini protein gel, wet transferred onto a nitrocellulose (NC) membrane at 30 V for 4 hours at 4 ℃, and subjected to chemiluminescent biotinylated nucleic acid detection. The same NC membrane was finally subjected to Western blot for TLR7 detection using rabbit anti-human TLR7 (1:1000) in combination with horse radish peroxidase (HRP)-conjugated goat-anti-ribbit secondary antibody (1:5000).

### Proximity biotinylation of csRNA-proximal proteome

The experiment was performed using three independent cell cultures for each treatment. Cells in a 15 cm dish at 90% confluency were washed twice with 1x DPBS after removing culture media, and was lifted with 3 mL per dish 10 mM EDTA in 1x DPBS. For csRNA-digested sample, RNase A was added to the lifting buffer at a final concentration of 40 μg/mL. 1x DPBS was used to bring cell suspension to 10 mL, which was transferred to a 15 mL Falcon tube. The dish was washed with 5 mL 1x DPBS, which was combined in the same tube. The cells were spun down at 300 g for 5 minutes at 4℃, and pellets were incubated for 30 minutes at 4 ℃ with per dish 250 μL solution of precomplexed TLR7-Fc (5 μg/mL final concentration) and protein-A-HRP conjugate (Thermo Fisher Scientific, 101023, 7 μg/mL final concentration) in 0.5% BSA in 1x DPBS containing 200U/mL RNasein. For an IgG isotype control, human IgG (Thermo Fisher Scientific, 02-7102) was premixed with protein-A-HRP at the same concentration. Cells were washed three times with cold 1x DPBS, and spun down at 200 g for 2 minutes at 4 ℃ each time. After the final wash, the pellets were resuspended in 1 mL per sample 500 μM biotin-phenol in 1x DPBS, and let sit for 1 minute at room temperature. 1 mL 2 mM hydrogen peroxide in 1x DPBS was added to each sample and mixed well. This mixture was incubated for 2 minutes at room temperature and 10 mL cold quenching buffer (5 mM Trolox + 10 mM sodium ascorbate + 10 mM sodium azide in 1x DPBS) was directly added to the sample. The cells were washed two more times with quenching buffer, and were lysed in 300 μL per sample RIPA buffer (50 Mm Tris pH 7.5, 150 mM NaCl, 0.5% sodium deoxycholate, 0.1% SDS, 1% NP40) containing 1x cOmplete™ protease inhibitor cocktail and 1mM PMSF. The lysate was sonicated to clarity, and was spun down at 21,000 g for 10 minutes at 4℃. An aliquot of the supernatant was taken for WB to detect biotin signal, and the rest was subjected to enrichment of biotinylated proteins.

The affinity pulldown follows the previous report by Santos-Barriopedro et al. The pellets were taken up in RIPA buffer. The supernatant from the sonicated lysates was subjected to bead enrichment with 12.5 μL per sample streptavidin sepharose high-performance beads (Cytiva, 15511301) on an end-to-end rotator for 2 h at room temperature. The beads were washed 5 times with 500 μL RIPA buffer and 4 times with 500 μL PBS, prior to LC-MS/MS sample preparation.

### Live-cell surface biotinylation, UVC crosslinking followed by OOPS

Cells in a 15 cm dish were cultured to 85 – 90% confluency. Prior to detachment, culture medium was removed and the cells were washed once with 1x DPBS. The cells were lifted using 10 mM EDTA in 1x DPBS and transferred into 15 ml Falcon tubes. The pellets were then taken up in 1 ml per tube biotinylation solution containing 2.5 mM sulfo-NHS-SS-biotin (Thermo Fisher Scientific, 21331). For non-biotinylated control, cells were treated with just 1x DPBS and were processed the same way. The cells were incubated at room temperature for 15 min with gentle shaking every 5 min. Biotinylation was quenched using another 8 ml per tube 20mM Tris-HCl in 1x DPBS pH 7.4 at room temperature for 10 min. After spin-down, the cells were taken up in 7 ml 1x DPBS and transferred into a 10 cm dish for UVC crosslinking following procedures described above. The crosslinked cells were collected by centrifugation and lysed in TRIzol for performing OOPS. The OOPS procedure is described in detail in Section *UVC and eosin Y-mediated crosslinking and orthogonal organic phase separation (OOPS)* below.

The RNA-bound RBP from organic phase was re-dissolved in cell lysis buffer (50mM pH7.5 Tris-HCl, 100mM NaCl, 1% (v/v) NP-40, 0.1% (v/v) SDS, 0.5% (w/v) Sodium deoxycholate), and subjected to affinity purification using agarose beads functionalized with immobilized neutravidin (Thermo Fisher Scientific, 29204). The bead binding was performed at 4°C for 2 hours on an end-to-end rotator. The supernatant post-enrichment was discarded. The bead was washed 5 times with lysis buffer, and a final wash in 1x DPBS, prior to LC-MS/MS sample preparation.

### Proteomics sample preparation and LC-MS/MS analysis

Beads from biotin affinity pulldown were incubated with 50 μL elution buffer (2 M Urea, 10 mM DTT, 100 mM Tris pH 8) on a thermoshaker at 1250 rpm for 20 minutes at room temperature. Iodoacetamide (50 mM final concentration) was then added, and the samples were incubated in a thermoshaker at 1250 rpm in dark for 10 minutes at room temperature. 2.5 μL of trypsin (0.1 mg/mL stock solution) was then added, followed by incubation in a thermoshaker at 1250 rpm for 2 hours at room temperature. Samples were spun down (1500 x *g*, 2 min, RT) and supernatant was collected. Another 50 μL elution buffer was then added to the beads, which were incubated on a thermoshaker at 1250 rpm for 5 minutes at room temperature. Beads were then spun down (1500 x *g*, 2 min, RT), and the supernatant collected and combined with the first elution. 1 μL trypsin (0.1 mg/mL) was added and the samples were incubated overnight at room temperature. Tryptic peptides were acidified by adding 10 μL 10% (v/v) TFA, and purified using C18 Stagetips^85^ (3M Empore). Briefly, StageTips were prepared and washed with methanol followed by a wash with buffer B (80% acetonitrile and 0.1% formic acid) and two washes with buffer A (0.1% formic acid). All washes were done at 1500 × *g* for 4 min at RT. Samples were loaded onto the StageTips and centrifuged at 600 × *g* for 10 min at RT and washed once with buffer A. StageTips were stored at 4°C

Peptides were eluted from StageTips with 30 ml of buffer B, reduced to a volume of 5 ml by Speedvac centrifugation, after which 7 ml of buffer A (0.1% formic acid) was added. For the TLF7 proximity labeling experiments, 6 µl of each sample was loaded on an Easy LC1000 (Thermo Scientific) connected to an Orbitrap Exploris 480 (Thermo Scientific). The machine was operated in data-dependent acquisition (DDA) mode, with acquisition time set at 60 minutes. An acetonitrile gradient of 12-30% in 43 minutes was used followed by an increase of the acetonitrile concentration to 60% in 10 minutes and to 95% in 1 minute. The full scan of the peptides was set to a resolution of 120,000 in a scan range of 350–1300m/z. For biotinylation-crosslinking-OOPS experiments, 3 µl of each sample was loaded on a Vanquish Neo UHPLC connected to a Orbitrap Astral. The machine was operated in data-independent acquisition (DIA) mode, with acquisition time set at 24 minutes. An acetonitrile gradient of 8-35% in 15 minutes was used followed by an increase of the acetonitrile concentration to 45% in 3 minutes and to 99% in 30 seconds. The full scan of the peptides was set to a resolution of 240,000 in a scan range of 150–2000 m/z.

### Proteomics data analysis

Raw mass spectrometry data were analysed using Proteome Discoverer version 3.1 (Thermo Scientific) and searched against the human UniProt/SwissProt database.

For DDA, we used the built-in processing workflow “PWF_OT_Precursor_Quan_and_LFQ_CID_SequestHT_Percolator” and the built-in consensus workflow “CWF_Comprehensive_Enhanced Annotation_LFQ_and_Precursor_Quan”, with default settings. For the Sequest HT search, database parameters were enzymatic digestion with trypsin allowing two missed cleavages, a minimum peptide length of 6 amino acids and a maximum peptide length of 144 amino acids. We used a precursor mass tolerance of 10 ppm and a fragment mass tolerance of 0.6 Da. Cysteine carbamidomethylation was included as a static modification (57.021 Da), while methionine oxidation (15.995 Da) and protein N-terminal acetylation (42.011 Da) were included as dynamic modifications. FDR filtering was performed via percolator with a strict target FDR of 0.01 and a relaxed FDR of 0.05. Strict parsimony was applied for protein grouping, and unique plus razor peptides were used for quantification. Peptide quantification normalization was applied based on total peptide amount. Imputation Mode was set to “Low Abundance Resampling”. “Proteins” tab was exported for downstream processing and statistical analysis with R.

For DIA, we used the built-in processing workflow “PWF_Astral_CHIMERYSonArdia_DIA_MBR.BestPSM” and the built-in consensus workflow “CWF_Comprehensive_Enhanced Annotation_LFQ_and_Precursor_Quan”, with default settings. For the CHIMERYS search, database parameters were enzymatic digestion with trypsin allowing two missed cleavages, a minimum peptide length of 7 amino acids and a maximum peptide length of 30 amino acids. We used a fragment mass tolerance of 20 ppm. Cysteine carbamidomethylation was included as a static modification, while methionine oxidation was included as dynamic modification. FDR filtering was performed via percolator with a strict target FDR of 0.01 and a relaxed FDR of 0.05. Strict parsimony was applied for protein grouping, and unique plus razor peptides were used for quantification. Peptide quantification normalization was applied based on total peptide amount. Imputation Mode was set to “Low Abundance Resampling”. “Proteins” tab was exported for downstream processing and statistical analysis with R.

For downstream processing, proteins were first filtered for having at least 2 unique peptides. Proteins containing “KRT” in their names as well as proteins with less than 3 values in at least 1 condition were filtered out. Statistical analysis was done using the DEP package, which uses a Student’s t-test. Criteria for enrichment are listed in the figure legends. Volcano plots were generated by plotting the –log10 of the p-value against the log2 fold change. Gene ontology term searches for identified hits were performed using RBP2GO.

The proteomics datasets have been deposited in the ProteomeXchange Consortium via the PRIDE^86^ partner repository with the dataset identifiers PXD061693. (Reviewer log-in username. Password: ncGUlhn5XtSB)

### Genetic screening and data analysis

For knockout screening, the human CRISPR Brunello genome-wide knockout library was a gift from David Root and John Doench (Addgene 73178). For virus production, HEK 293T cells were transfected with packaging plasmids pRSVrev, pHCMV-G VSV-G and pMDLg/pRRE together with the Brunello plasmid using polyethyleneimine (Polyscience Inc.). Virus was harvested, filtered and 150 million Mel526 melanoma cells (a kind gift from Ramon Arens) were transduced in the presence of 8 µg/ml polybrene (Millipore) at an MOI of 0.3. Transduced cells were selected using puromycin (1 µg/ml) and after seven days, two batches of cells were stained for surface RNA using TLR7-Fc and the lowest 5% of expressing cells were sorted. Cells were grown out and sorted again using the same gating strategy as for the first sort. After this sort, genomic DNA was isolated for both the unsorted and sorted populations and gDNAs were amplified using the established protocol.^87^ gRNAs were sequenced using the Illumina NovaSeq6000 and inserts were mapped to the reference. Analysis of gRNA enrichment was done using PinAPL-Py.^88^

### Genetic screening hit validation

Hits from the knockout screen were validated using the top scoring gRNAs of each gene, which were cloned into the lentiCRISPR v2 vector (a gift from Feng Zhang, Addgene plasmid no. 52961). Mel526 cells were stably transduced with these gRNAs, selected using puromycin, and pooled knockout cells were used for the analysis. Guide sequences were as follows: EXT2, 5′-CACCGTGGTTAAGCACATCGATGGA-3′; EXTL3, 5′-CACCGAAATGAACCTCGGTAACACG-3′; HS6ST1, 5’-AAACCCGCGAGACTTGGCTCTTCTC-3’, C3orf58, 5’-

CACCGGCAGACGCACGTCGCCGTTG-3’. Mutant cells were subjected to flow cytometry analysis to check for TLR7-Fc binding, following the procedure described above.

### Preparation of PVR KO mutant cells

HEK293T cells (ATCC CRL-3216) were seeded into a 10 cm dish per plasmid at a confluency of 60-70% for lentiviral packaging. The utilized guide RNA was 5’-ATTGGTGCCCTCAAGCCAGG-3’ (PVR, NM_006505.4, NC_000019.10. Sense, exon. Position: 44650019). For a 10 cm dish, 5.63 µg pCMV-VSVG, 8.55 µg pMDLg-RRE (gag/pol), 4.5 µg pRSV-REV and 11 µg pRRL were combined with serum-free DMEM to a total volume of 500 µl. In a second tube, 90 µg of PEI for a 10 cm dish was mixed with serum-free medium to a total volume of 500 µl. This mixture (tube two) was then slowly added to tube one and thoroughly mixed. After 15-30 minutes, the resulting mixture was added dropwise to the cells after adding 4 ml of serum-free medium. 4 hours post-transfection, 4 ml of DMEM with 20% FCS was supplemented to the plate. Subsequently, the plates were incubated overnight at 37°C with 5% CO_2_. 48 hours post-transfection, the supernatant containing virus was harvested by collecting the medium into a sterile 15 ml tube. The collected medium centrifugated at 3000 rpm at 4°C for 10 minutes and was subsequently filtered through a 0.45 µm low protein binding membrane (Millipore). The viruses were utilized immediately or stored at −80°C. The transduced cells were cultured using high glucose DMEM with 10% FBS and 1% Pen-Strep. Transduction was executed by adding 30% of lentivirus plus 5 µg/mL Polybrene. Transduced cells were selected using puromycin (3 µg/ml) 48 hrs after transduction.

### UVC and eosin Y-mediated crosslinking and orthogonal organic phase separation (OOPS)

For suspension cells such as K562 (used in this study), start with ∼20 million. Culture media was removed and the cells were wash twice with 10 ml DPBS. Cell pellets were resuspended in 10 ml DPBS, and transferred into a 10 cm dish. For UVC crosslinking, the cells (without lid) were then UV-crosslinked with a Stratalinker 1800 chamber at 254 nm for 5 min on ice. The crosslinked pellets were collected and directly lysed in 1 ml TRIzol per million cells. For EY-mediated crosslinking, cells after wash were resuspended in DPBS containing 50 µM eosin Y (CAS #17372-87-1). The cells were placed in dark at room temperature for 2 min and put on ice with lid open. The cells were exposed to green light (100 W chip-on-board) for 10 min. The crosslinked pellets were collected and directly lysed in 1 ml TRIzol per million cells.

For adherent cells, start at least with a 10 cm dish. Culture media was removed and the cells were wash twice with 10 ml HBSS. In case of UVC crosslinking, upon removal of the wash solution, the dish without the lid was placed in the chamber on ice, and exposed to UV for 5 min. The crosslinked cells were lysed in 1 ml TRIzol per 10 cm dish, and scraped to one side of the dish and transferred to a 2 ml Eppendorf tube. In case of EY-mediated crosslinking, 7 ml HBSS containing 50 µM eosin Y was added to the dish. The dish without lid was exposed to green light for 10min and washed once more with HBSS, followed by cell lysis in 1 ml TRIzol per 10 cm dish, scraping and the transfer as mentioned ealier.

OOPS was performed following reported literature. Unless otherwise noted, the phase separation procedure and the amounts of enzymes and duration of incubation for releasing RNA-bound proteins and protein-bound RNA remain the same as reported. In brief, 0.2 ml chloroform per 1 ml TRIzol was added to the tube, followed by vortexing and spin-down at 4°C. The aqueous and organic phases were carefully removed with a blunt needle without touching the interphase. Such procedure was repeated 3 times for UVC crosslinking and 4 times for EY crosslinking. The cleaned interphase was then precipitated with methanol. The resulted pellets were wash once again with methanol and air-dried for 5 min.

To isolate protein-bound RNA, the interphase pellet was directly treated with proteinase K solution. The protein-degraded RNA samples were then purified with TRIzol in combination of Zymo RNA Clean & Concentrator columns (Zymo Research C1008) following the product manual. The cleaned RNA samples were then subjected to 1% agarose gel electrophoresis and stained with SybrGold. The stained gel was imaged with a ChemiDoc system using SybrGold settings.

To isolate RNA-bound proteins, the interphase pellet was redissolved in 1% SDS containing ammonium bicarbonate. The RNA was fragmented by sonication and Mg(II) ions at 94°C. To the resulted solution, RNase Cocktail was added. The digestion was incubated at 37°C for 20 hours, followed by another round of TRIzol phase separation. The RNA-bound proteins were collected from the organic phase and precipitated with methanol, redissolved in 1% SDS. An equal volume of each sample solution was taken for SDS-PAGE and Western blot. To check for total isolated proteins, the gel was Coomassie-stained and scanned under a ChemiDoc system. For Western blot, the PVDF membrane post-transfer was blocked with 5% BSA in TBST (blocking buffer), incubated with the corresponding primary antibody (1:1000 dilution for all) in blocking buffer, washed 3 times with TBST, and incubated with the secondary antibody (1:5000 – 1:10000) in blocking buffer. The resulted membrane was washed 3 times, treated with West Pico PLUS Chemiluminescent Substrate (Thermo Fisher Scientific, 34580), and imaged under a ChemiDoc system.

### Conjugation of eosin isothiocyanate to streptavidin

Eosin 5-isothiocyanate (Abcam, ab270343) was dissolved in DMSO to make a 100mM stock solution. The conjugation reaction was performed for 500 µg streptavidin in 0.5 ml 1X DPBS at pH 8, with eosin isothiocyanate at a final concentration of 2mM. The reaction was allowed to proceed for 2 hours at room temperature, with shaking and in dark. The resulting mixture was purified twice with MidiTrap G-25 (GE Life Sciences, 28922530) columns, and buffer exchanged into 1x DPBS pH 7.4. The purified conjugate was stored at 4 °C in dark.

### Enzyme-linked immunosorbent assay (ELISA) of EY-conjugated streptavidin

The 96-well MaxiSorp microtiter plate was coated with either streptavidin-EY or unmodified streptavidin at 10 µg/ml in 100mM NaHCO_3_ overnight in dark at room temperature. The wells were washed twice with PBS and then blocked with 2% BSA in PBST (blocking buffer) 1 hour in dark at room temperature. Serial dilutions (3x, starting concentration 10 µg/ml) of a biotinylated mouse monoclonal antibody (BioLegend, 317320) in blocking buffer were added to each row, and was incubated for 1 hour in dark at room temperature, followed by washing 3 times with PBST. To each well, HRP conjugated alpaca-anti-mouse (Jackson ImmunoResearch, 615-035-214) in blocking buffer was added and allowed to incubate for 45 minutes in dark at room temperature, followed by washing 3 times with PBST. Finally, 1-step Ultra TMB-ELISA (Thermo Fisher Scientific, YJ4085531) and incubated for 2 mins, and was stopped by 2M sulfuric acid. The optical density at 450 nm (OD450) was measured by a microtiter plate reader.

### Spatio-selective crosslinking followed by OOPS (SCOOPS)

Cells at 85-90% of confluency in a 10 cm dish was used for each sample. For csRNA degradation, cells were treated with 13 µl of RNase A (Thermo Fisher Scientific, 12091021) and 3.2 µl of RNase I (Jena Biosciences) in 6.5 mL culture media under standard culturing condition (37°C, 5% CO_2_ and humidified atmosphere) for 30 mins. All cells were washed with HBSS, and were incubated with 6 ml culture media containing 5 µg/ml biotinylated WGA (Vector Laboratories, B-1025) at 4°C for 1 hour. The corresponding negative control was with merely 6 ml culture media without biotinylated WGA. After aspiration, cells were washed twice with HBSS and then incubated with 6ml of culture media containing 2.5 µg/ml streptavidin-EY at 4°C for 30 mins in dark, after which the cells were washed twice with HBSS. The corresponding negative controls was with unmodified streptavidin in the culture media. 6mL HBSS was added to each plate prior to green light irradiation at 4°C for 10 mins. Supernatant was immediately removed after irradiation and the cells were lysed in 1.4ml TRIzol per dish, scraped and transferred into a 2 ml Eppendorf tube for OOPS. The OOPS procedure is the same as described earlier. Protein-bound RNA was released from interphase, purified, and subjected 3’-end Cy5 ligation, which followed the same procedure for 3’-end biotin ligation described above, except for the purification was performed using Zymo RCC-5 columns. 2% (v/v) of the first aqueous phase containing total cellular RNA was taken as loading control, and was also ligated with Cy5.

### RNA sequencing library preparation

The RNA sequencing library preparation procedures are adapted from what was described for ‘size-matched input’ in the published reports.^29^ All library preparations started with 150 ng (Qubit RNA quantification, broad range) RNA input. In brief, the RNA was first subjected to 3’-end repair (total volume 20 µl per sample): 2 µl 10x T4 PNK buffer, 2 µl FastAP, 1 µl T4 PNK, 0.5 µl SuperRasIn, 0.5 µl Turbo DNase, 37 °C, 45 min, 900 rpm on a thermoshaker. The end-repaired RNA was purified using TRIzol in combination with Zymo RCC-5 columns following the product manual, and eluted in 10 µl nuclease-free water. The pre-adenylated 3’-adapter was ligated using the following procedure (total volume 20 µl per sample): 2 µl 10x RNA ligase buffer, 10 µM (final concentration) pre-adenylated adapter, 6 µl 50% PEG8000, 0.3 µl DMSO, 0.5 µl SuperRasIn, 2 µl T4 RNA ligase 1 high concentration, 16 °C, 3h, 1000 rpm on a thermoshaker. The excess adapter was degraded post-ligation using 1 µl 5′ deadenylase (NEB, M0331S) and 2 µl RecJf (NEB, M0264S) after adding 2.5 µl 10x NEB buffer 1. The mixture was purified using TRIzol in combination with Zymo RCC-5 columns, and eluted with 10 µl nuclease-free water. Reverse transcription was performed with the following protocol: 4 µl 5x first strand buffer, 10 mM DTT, 0.8 mM dNTP, 0.5 µl SuperRaseIn, 1 µl SuperScript III (Thermo Fisher Scientific, 18080093), 55 °C, 30 min, on PCR heat block. The excess primer was degraded using 3 µl ExoSAP-IT, quenched with 1 µl 0.5 M EDTA. RNA was then removed under alkaline pH (3 µl 1 M NaOH) at elevated temperature (70 °C, 12 min) and neutralized with 3 µl 1 M HCl. The resulted complementary DNA (cDNA) was purified using MyOne Silane beads, and eluted with 20 µl nuclease-free water. Half of the purified cDNA was subjected to PCR amplification with NEBNext Multiplex Oligos for Illumina: 30 µl total reaction volume, 2x KAPA HiFi HotStart ReadyMix (Roche, 07958927001), 0.3 µM forward and reverse primers. Initial denaturation, 95 °C, 3 min; denaturation, 98 °C, 20 s; annealing, 65 °C, 15 s; extension, 72 °C, 15 s; total cycle number, 16; final extension, 72 °C, 1 min. The amplified library was first purified with AmpureXP beads (Beckman Coulter, A63880) and then size-selected using PAGE gel electrophoresis. The resulted libraries were quantified on a Qubit tube reader and Bioanalyzer and finally mixed at equal molarity to make the 20 nM pooled library for Illumina sequencing on a Illumina Novaseq X plus platform at BMKGENE GmbH.

### Sequencing data analysis

UMIs (NNNNN) were extracted from the demultiplexed pair-end fastq files and appended to read identifiers by UMI-tools.^89^ The adapters were trimmed by Cutadapt.^90^ Read1 adapter to trim, AGATCGGAAGAGCACACGTCTGAACTCCAGTCAC; Read2 adapter to trim, GATCGTCGGACTGTAGAACTCTGAACGTGTAGATCTCGGTGGTCGCCGTATCATT. The processed reads were pseudo-aligned and quantified at transcript level using Kallisto quant,^91^ due to its outstanding ability to probabilistically assign multi-mapping reads to transcripts at high accuracy. The pseudo-alignment was done against the reference transcriptome of Gencode Release 47. The read counts for individual transcripts were normalized as RPMs.

### KIR2DL5 transfection, in-situ crosslinking and RNA co-precipitation

Wildtype (CHO-K1) and heparan sulfate-deficient cells (CHO pgsD-677) in a 10 cm dish at 70% confluency were transfected with 10 μg full-length KIR2DL5 plasmid (Addgene, #157623), using 2μl polyetherimide (PEI, 2mg/ml) per μg DNA in F-12K medium. 6h after transfection, the serum-free medium was replaced the one supplemented with 10% fetal bovine serum (FBS). The next day, cells were washed once with 1x DPBS following culture media removal. The cells were crosslinked in a Stratalinker for 5 min with the lid open. Once crosslinking is finished, the cells were scraped into 500μl per dish lysis buffer (50 mM Tris-HCl pH 7.4, 100 mM NaCl, 1% v/v NP-40, 0.1% v/v SDS and 0.5% w/v sodium deoxycholate) containing 1x cOmplete™ protease inhibitor cocktail (Roche, 11836153001). The cells were transferred into an Eppendorf tube and sonicated for 5 min (20% power, 5s on and 5s off). To the sonicated lysate was added 2 μl Turbo DNase, and 4 μl diluted RNase I (prediluted 1:20 in 1x DPBS). The lysate was then incubated at 37 °C for 5 minutes, followed by the addition of 2.5 μl SUPERaseIn. After centrifuging at 15,000g for 10 min at 4 °C, the supernatant was transferred to antibody-bound protein A beads (rabbit anti-DYKDDDDK tag antibody, Proteintech, 20543-1-AP; or rabbit IgG isotype control). The suspension was incubated on an end-to-end rotator at 4 °C for 2 h. The supernatant was magnetically separated and the beads were washed twice with 200 μl high-salt wash buffer (50 mM Tris-HCl pH 7.4, 1 M NaCl, 1% v/v NP-40, 1 mM EDTA, 0.1% v/v SDS and 0.5% w/v sodium deoxycholate), three times with 200 μl low-salt wash buffer (20 mM Tris-HCl pH 7.4, 10 mM MgCl_2_, 0.2% v/v Tween-20 and 5 mM NaCl). The precipitated RNA-KIR2DL5 was 3’-end repaired and biotin ligated following the procedure described for TLR7-Fc. After the final wash, beads were directly taken up in 1x SDS sample loading buffer containing 1x sample reducing solution, denatured at 75°C for 10 minutes. The samples were resolved with SDS-PAGE, wet transferred onto a nitrocellulose (NC) membrane at 30 V for 4 hours at 4 °C, and subjected to chemiluminescent biotinylated nucleic acid detection using streptavidin-HRP (Thermo Fisher Scientific, 89880D). For Flag tag detection, the samples were resolved with SDS-PAGE, semi-dry transferred onto a polyvinylidene fluoride (PVDF) membrane at 25 V for 30 mins, and detected using rabbit anti-DYKDDDDK tag (Proteintech, 20543-1-AP, 1:10000) in combination with HRP-conjugated goat-anti-ribbit secondary antibody.

### Biotinylation at total RNA fragments 3’-end (for in-vitro KIR2DL5 binding)

Isolated total RNA was treated with Turbo DNase and then proteinase K to remove residual DNA and polypeptides. The purified RNA was fragmented with 4 μl 10x RNA Fragmentation Buffer (NEB, E6186AVIAL) added to 36 μl of total RNA in DEPC-treated water. The reaction was incubated at 94 °C for 2.5 minutes. Afterwards, 5 μl RNA Stop Solution (NEB, E6187AVIAL) was added. The fragmented RNA was kept on ice for over a minute was purified with the Zymo RCC-25 columns following the protocol provided by the supplier. To perfom end repair, 80 μg purified fragmented RNA was combined with 10 μl 10x PNK buffer, 5 μl FastAP (Thermo Fisher Scientific, EF0654), 1 μL SUPERase·In and 2 μl Turbo DNase, and incuabted at 37 °C for 30 minutes on a thermoshaker. The dephosphorylated RNA was combined with 1 ml TRIzol, from which the aqueous phase was mixed with an equivolume of ethanol and purified using Zymo RCC-25 columns. The RNA was eluted in 50 μl nuclease-free water. To the RNA solution, 14 μl 5x polymerase buffer (100 mM Tris-HCl pH 7.0, 3 mM MnCl_2_, 0.1 mM EDTA, 1 mM DTT, 0.5 mg/ml acetylated BSA, 50% glycerol (v/v)), 5 μl yeast poly(A) polymerase (Jena Bioscience, RNT-006-S), 2’-Azido-2’-dATP (Jena Biosciences, NU-976S, 50 μM final concentration) and 0.5 μl SUPERase·In were added. The solution was incubated at 37 °C for 1 hour on a thermoshaker. The resulted RNA-3’-azide was purified as described for the end repair. Finally, 50 μl RNA-3’-azide was combined with 20 μl of 2x denaturing loading buffer (95% formamide, 25 mM EDTA). Next, the RNA was incubated with 1 mM of DBCO-biotin at 50 °C for 10 minutes. The biotinylated RNA was purified as described above.

### In-vitro KIR2DL5 binding, crosslinking, and RNA co-precipitation

The 3’-end biotinylated RNA described above was denatured in DEPC-treated water at 70 °C for 5 minutes and immediately put on ice. KIR2DL5-Fc (Biotechne, 6634-KR) or IgG isotype control (0.5 μg) were incubated with 2 μg biotinylated or non-biotinylated RNA in DPBS lysis buffer (1% NP-40 (v/v), 0.1% SDS (w/v), sodium deoxycholate 0.5% (w/v), 1x DPBS) at 16 °C with gentle shaking for 30 minutes. Afterwards, the samples were irradiated on ice with UVC (254nm) for 5 minutes. Next, the samples were resolved on a 12% SDS-PAGE gel, wet transferred onto a nitrocellulose (NC) membrane at 30 V for 4 hours at 4 °C and subjected to chemiluminescent detection of biotinylated nucleic acid. The same NC membrane was subjected to Western blot for human Fc detection using AlexaFulor488-conjugated goat-anti-human IgG.

### AlphaFold3 prediction of structural model for KIR2DL5 and potential interacting partners

The prediction was performed on AlphaFold Server (https://alphafoldserver.com/). The sequence of KIR2DL5 extracellular domain (aa 22 – 238) was used as input. To predict nucleic acid interactions, octanucleotide repeats (A_8_, U_8_, G_8_ and C_8_) were used as input. To check for potential protein-protein interactions with cell-surface RBPs, the sequences of overlapping hits between the TLR7-proximal and biotinylaiton-crosslinking-OOPS were obtained from Uniprot and used directly as another protein entity.

---

**Supplementary Figure 1.**
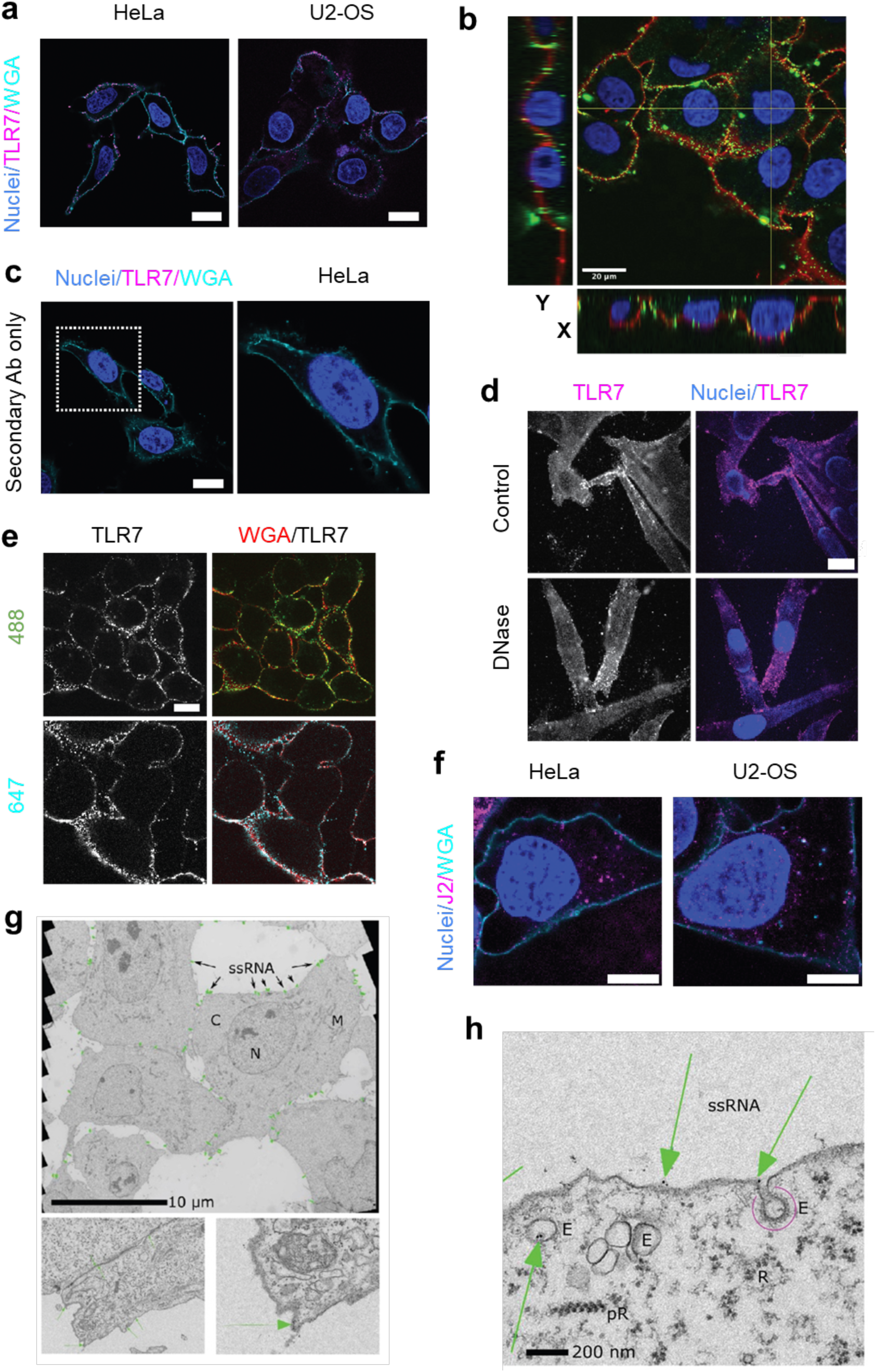
Immunofluorescence imaging on different cell lines using TLR7-Fc. **a**) csRNA staining on HeLa and U2-OS cells probed by TLR7-Fc and AlexaFluor647-conjugated rat-anti-human IgG Fc antibody (HeLa) or AlexaFluor488-conjugated goat-anti-human IgG Fc antibody (U2-OS) pseudocoloured as magenta. Cell surface was stained by WGA-biotin and streptavidin-AlexaFluor488 (HeLa) or streptavidin-AlexaFluor594 (U2-OS), pseudocoloured as cyan. **b**) Optical sections through U2-OS cells labeled with TLR7-Fc and AlexaFluor488-conjugated rat-anti-human IgG Fc antibody, pseudocoloured green. It shows the occurrence of csRNA on the cellular surface. Cell surface was stained by WGA-biotin and streptavidin-AlexaFluor594, pseudocoloured red. **c**) Representative confocal microscopy images of secondary antibody control (U2-OS cells) i.e. without TLR7-Fc but with AlexaFluor647-conjugated rat-anti-human IgG Fc antibody. Inset shows enlarged view. **d**) csRNA staining in the presence or absence of DNase I (10 U/ml) on Mel526 cells probed by TLR7-Fc and AlexaFluor647-conjugated rat-anti-human IgG Fc antibody. **e**) csRNA staining on living cells incubated at 4 ℃ with TLR7 and two different secondary antibodies in U2-OS cells (AlexaFluor488- or AlexaFluor647-conjugated anti-human IgG Fc antibodies. **f**) Confocal images of double-stranded RNA staining using HeLa and U2-OS cells probed by the J2 antibody and Cy3-conjugated donkey-anti-mouse IgG. Cell surface was stained by WGA-biotin and streptavidin-AlexaFluor488. Nuclear staining: DAPI. Scale bars: 10μm (**a**), 20μm (**b** - **f**). **g**) The ultrastructure of csRNA (green dots, indicated with arrows) localizations in U2-OS cells. TLR7- Fc served as a primary probe. 10nm gold nanoparticles conjugated to Protein A used as a secondary probe provide the necessary contrast in electron microscopy. Bottom panels provide different fields of view. **h**) Observation of csRNA internalized into the cell via endocytosis. The green arrows indicates the presence of csRNA inside endosomes with or without caveolae (pink arc). Abbreviations: C, cytosol; N, nucleus; M, mitochondria, E, endosome, R, ribosome; pR, polyribosome. Scale bars: 10μm (**g**), 200nm (**h**).

**Supplementary Figure 2.**
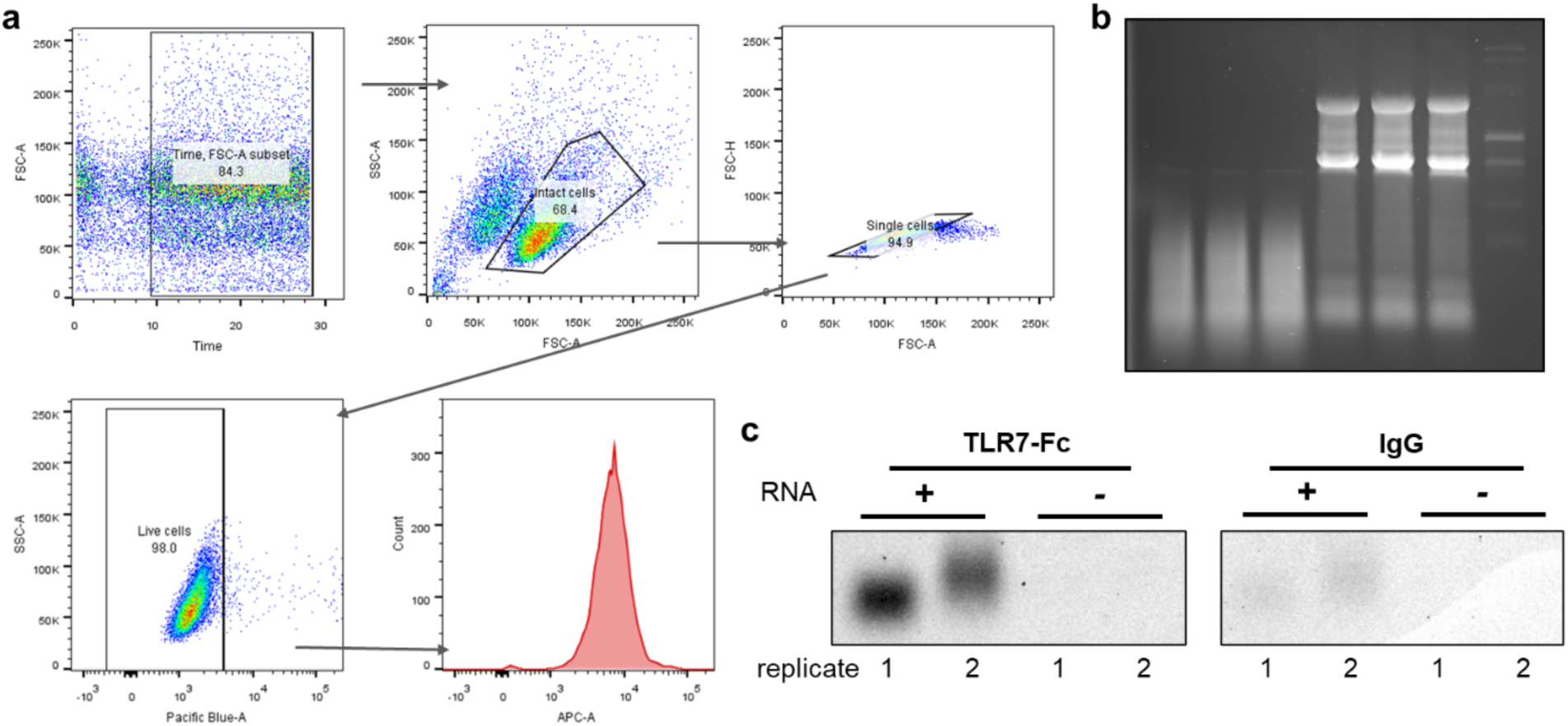
Additional data on flow cytometry analysis, exogenous RNA rescue, and in-vitro TLR7-RNA crosslinking and pull-down. **a**) Gating strategy for TLR7-Fc probed cells in flow cytometry. **b**) 1% agarose gel analysis of extracted total RNA. Left three lanes were 2-minute fragmented total RNA (from the RNA in the right three lanes) using magnesium-based fragmentation buffer. **c**) 1% agarose gel electrophoresis of on-bead 3’-end Cy5-labeled RNA released from bound TLR7- Fc. IgG was used as an RNA non-binding control. Replicate numbers mean the crosslinking experiments were performed with different batches of fragmented RNA independently. In replicate 1, 3’-end Cy5 ligation was performed overnight, and in replicate 2 for 4 hours.

**Supplementary Figure 3.**
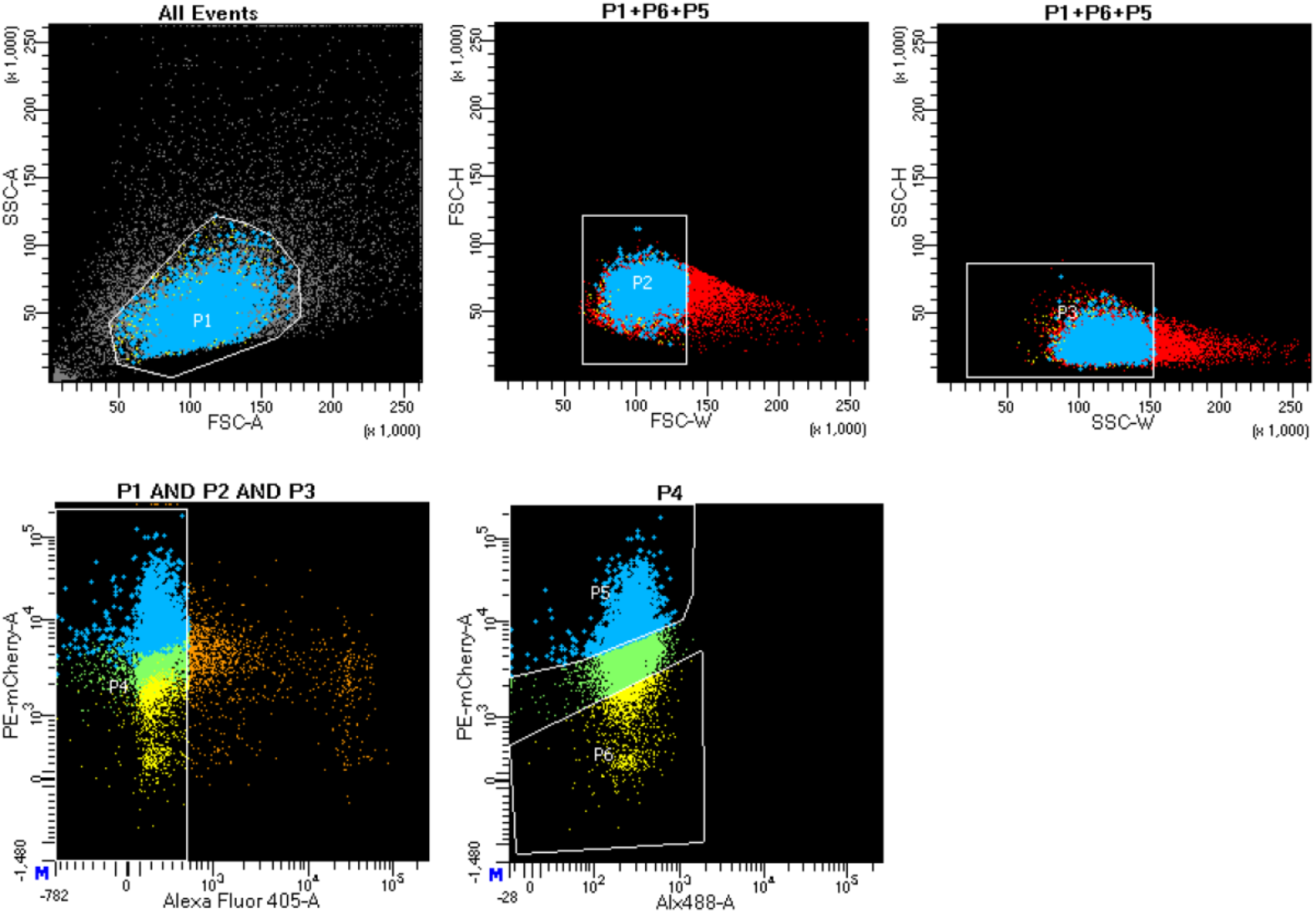
Gating strategies in FACS to enrich csRNA-low and -high phenotypes among lentiviral transduced cells. P1 gated for intact cells; P2 and P3 gated for singlets; P4 gated for living cells; P5 gated csRNA-high phenotype. P6 gated csRNA-low phenotype.

**Supplementary Figure 4.**
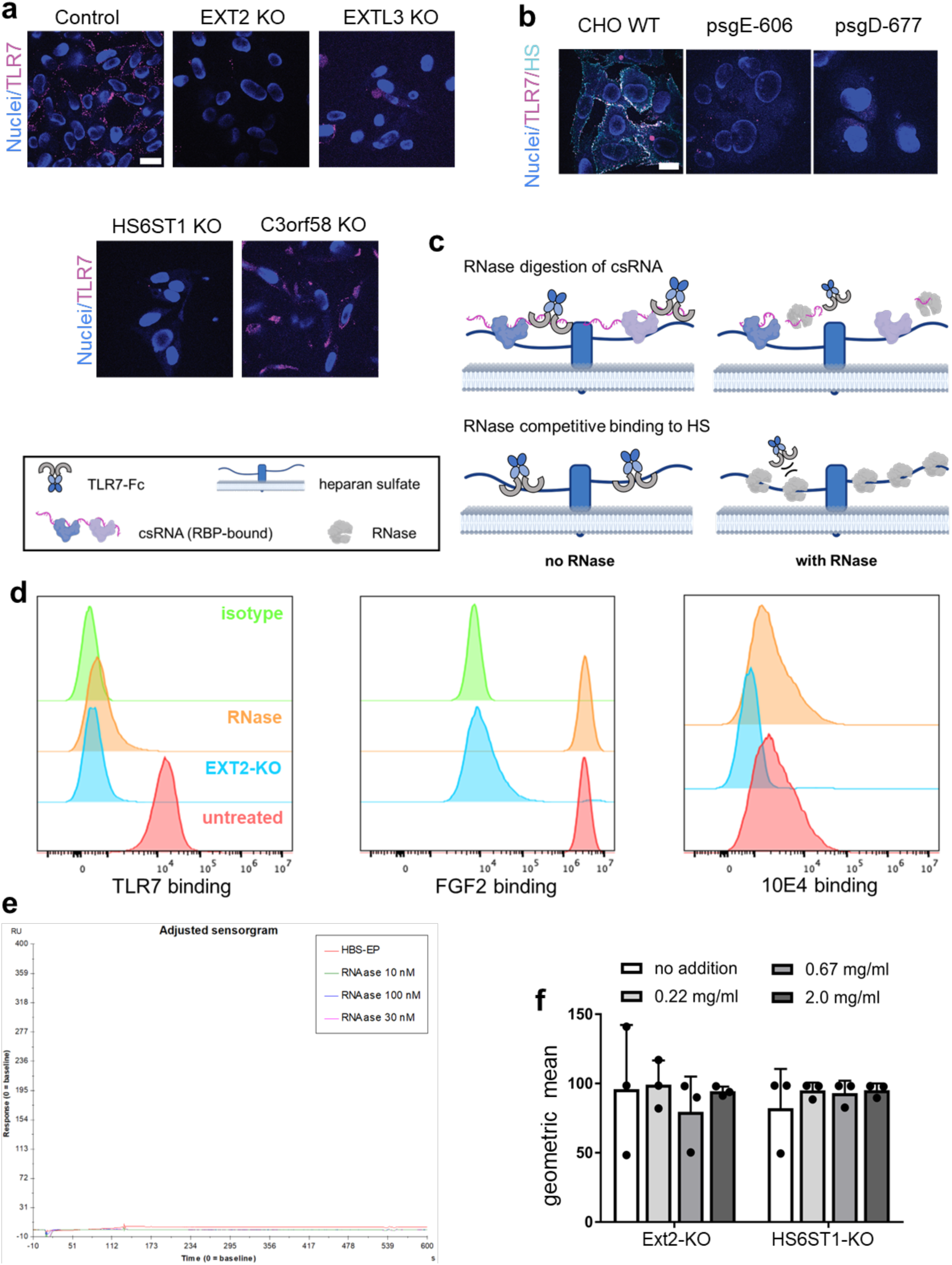
Validation of HS as essential factors for csRNA presentation. **a**) csRNA staining of EXT2, EXTL3, HS6ST1 and C3orf58 KO mutant cell lines (see Materials and Methods for sequences). TLR7-Fc was detected by AlexaFluor488-conjugated goat-anti-human IgG for csRNA (magenta). **b**) csRNA (magenta) and HS (cyan) staining of CHO-K1 (wild type) and heparan sulfate deficient cell lines. The primary secondary antibodies were the same as used in Figure 3d. Nuclear staining: DAPI. Scale bars: 20 μm (**d** - **e**). **c**) Schematic representation of an alternative scenario which could have given rise to the observation in genetic screening using TLR7-Fc. **d**) Histograms from flow cytometry analysis of RNase responsiveness of TLR7-Fc and well-known HS binders, including fibroblast growth factor 2 (FGF2) and anti-HS IgM (10E4 epitope). n = 3 independent cell cultures. Untagged recombinant FGF2 was pre-complexed with rabbit anti-FGF2. **e**) Surface plasmon resonance sensorgram of PureLink RNase A with heparin-coated chip. **f**) TLR7-Fc binding was not rescued by exogenous total RNA for HS-deficient mutant cells. n = 3 independent cell cultures. None of the groups is significant compared to no-RNA control (the white bar on the left). Two-tailed Student’s t-test, mean ± s.d.

**Supplementary Figure 5.**
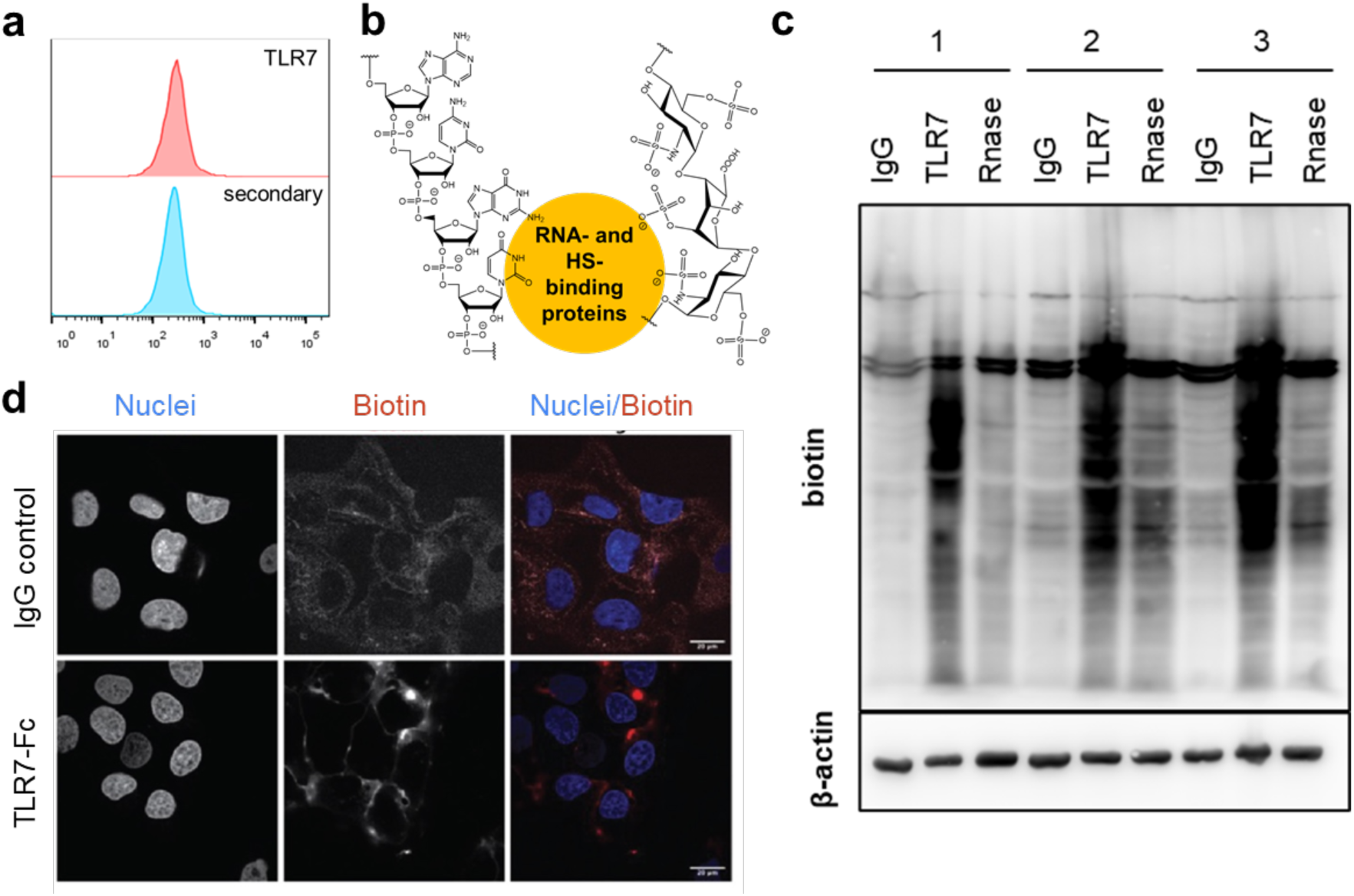
Rationales and supporting data for TLR7-mediated proximity labeling. **a**) Histograms from flow cytometry analysis of proteolytically lifted cells, which lost TLR7-Fc binding. **b**) Postulated model for RNA-HS association. Because both are negatively charged, they need a bridging molecule in between, which is hypothesized to be proteins. **c**) Western blot of cell-surface proximity biotinylated proteins. IgG, wildtype cells incubated with human IgG isotype control precomplexed with protein A-HRP; TLR7, wildtype cells incubated with TLR7-Fc precomplexed with protein A-HRP; RNase, extracellularly RNase A-treated cells incubated with TLR7-Fc precomplexed with protein A-HRP. The numbers above sample names indicate replicates. **d**) Fluorescence images of proximity biotinylation of extracellular RNase-treated cells. IgG was used as a control to check for background labeling. Red foci indicate biotinylation of HepRNA proximal proteins via biotin-phenol in the presence of HRP and hydrogen peroxide.

**Supplementary Figure 6.**
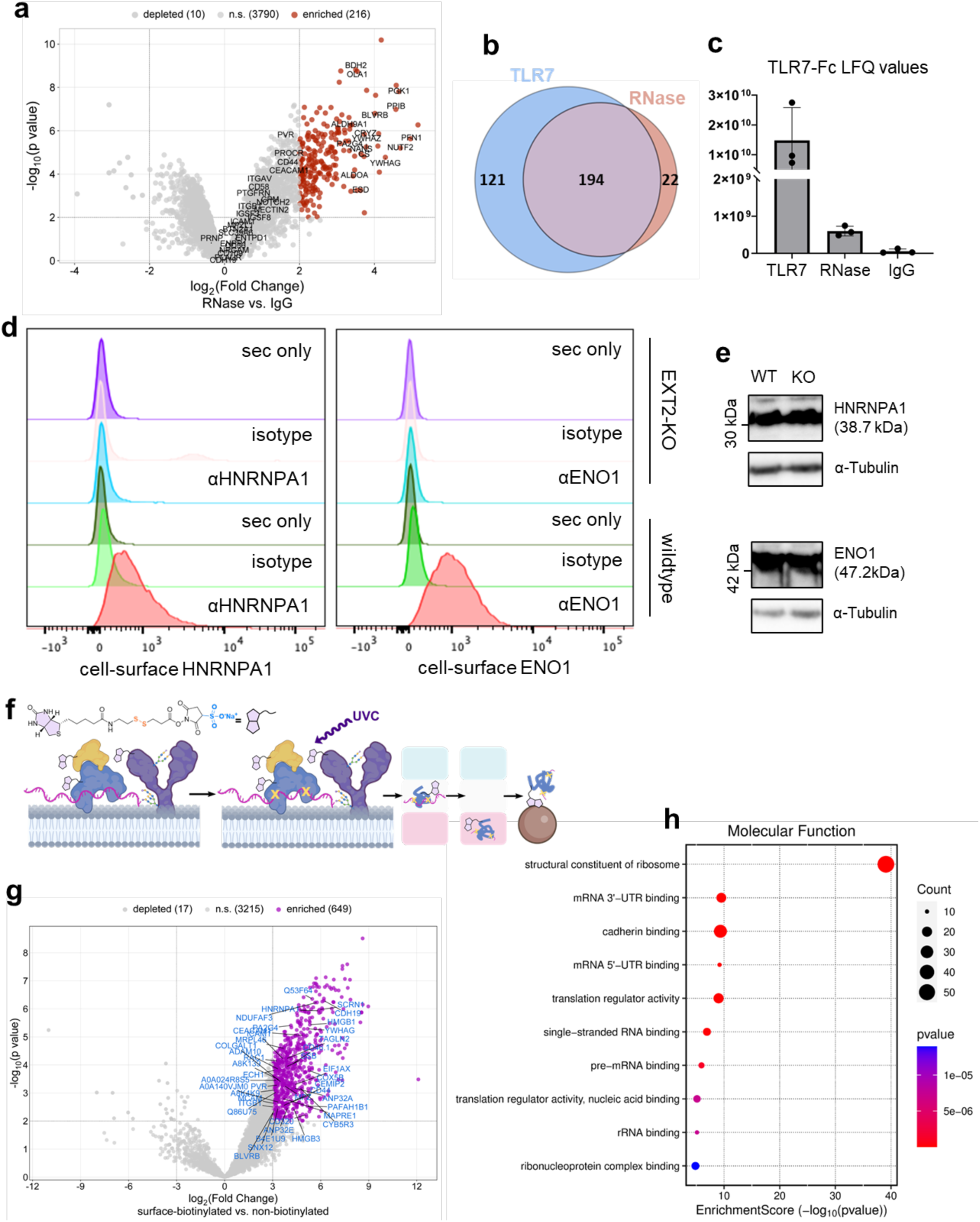
Validation of cell-surface localization and RNA complexation of csRBP candidates. **a**) Volcano plot from the comparison of proteome between TLR7-mediated proximity labeling post-RNase treatment and IgG control. hepRNA-proximal candidates are shown in red. Classically cell-surface localized proteins are labeled with their names to demonstrate loss of enrichment in RNase sample for many of these proteins. **b**) Venn diagrams showing consistently and uniquely enriched proteins (vs. IgG control) from TLR7 and RNase samples. **c**) Label-free quantification (LFQ) values of TLR7 indicates residual TLR7 were bound to cell surface after extracellular RNase treatment. **d**) Histograms from flow cytometry analysis demonstrating the presence of HNRNPA1 and ENO1 on the cell surface, and their dependency on intact HS chain. **e**) Western blot analysis of total cellular proteins indicates HS deficiency did not negatively affect the expression of HNRNPA1 or ENO1. **f**) Schematics of workflow for surface biotinylation, UVC crosslinking, OOPS and affinity pulldown to enrich csRNA-bound csRBPs. Cell-impermeable, amine-reactive biotin reagent tags the entire cell surface proteome. OOPS enriches RNA-bound proteins. **g**) Volcano plot from the biotinylation-OOPS experiment. The comparison is with biotinylated, OOPS-isolated samples over non-biotinylated, OOPS-isolated ones. Strongly enriched proteins (fold change ≥ 8, p < 0.01. n = 3 independent cell cultures) are shown in pink. csRBP candidates as TLR7-proximal proteins (**Supplementary Table 3**) are labelled with their names, and are considered to have been in complex with RNA on cell surface. **h**) Gene ontology analysis (molecular function) of strongly enriched proteins from biotinylation-OOPS experiments indicates the specificity of OOPS for isolating RNA-bound proteins.

**Supplementary Figure 7.**
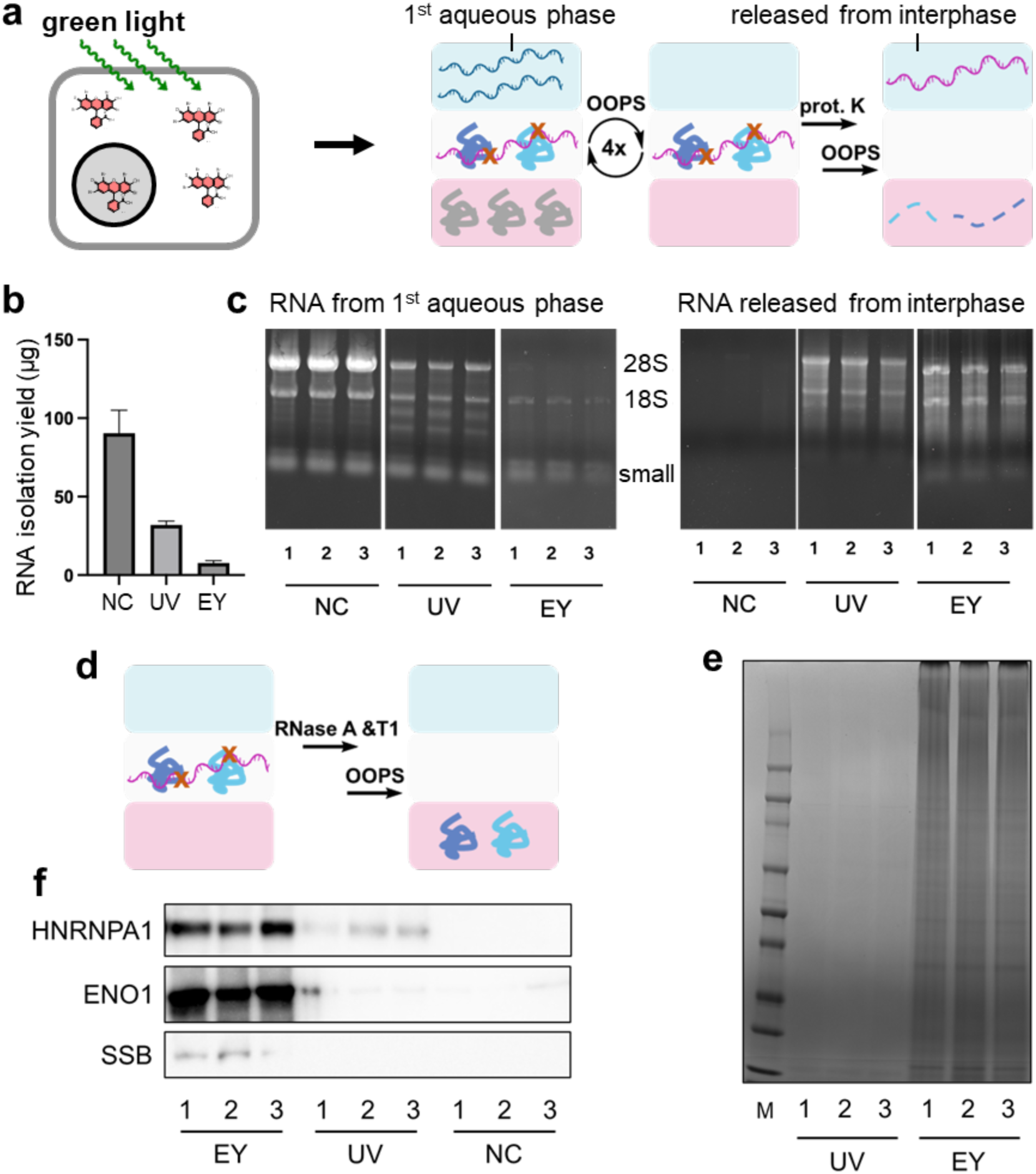
Observation of singlet oxygen-mediated RNA-protein crosslinking. **a**) Eosin Y (EY)-photosensitized singlet oxygen generation in living cells and APGC isolation of crosslinked RNA-protein complexes. Abbreviation: prot.K, proteinase K; OOPS, orthogonal organic phase separation. **b**) Yields of RNA isolated from the first aqueous phase decreased in EY- and UV-crosslinked cells compared to no-crosslinking (NC) control. n = 3 independent cell cultures. Error bars, s.d. **c**) 1% agarose gel electrophoresis of RNA isolated from the first aqueous phase and released from interphase after OOPS. Lane numbers in each sample represent independent cell cultures. **d**) Proteins crosslinked to RNA were released from the interphase and brought into the second organic phase. **e**) Coomassie staining of SDS-PAGE resolved proteins from the second organic phase of EY- and UV-crosslinked samples. **f**) Western blot of common RNA-binding proteins isolated in the second organic phase. HNRNPA1, heterogeneous nuclear ribonucleoprotein A1; ENO1, enolase 1; SSB, Sjögren syndrome antigen B.

**Supplementary Figure 8.**
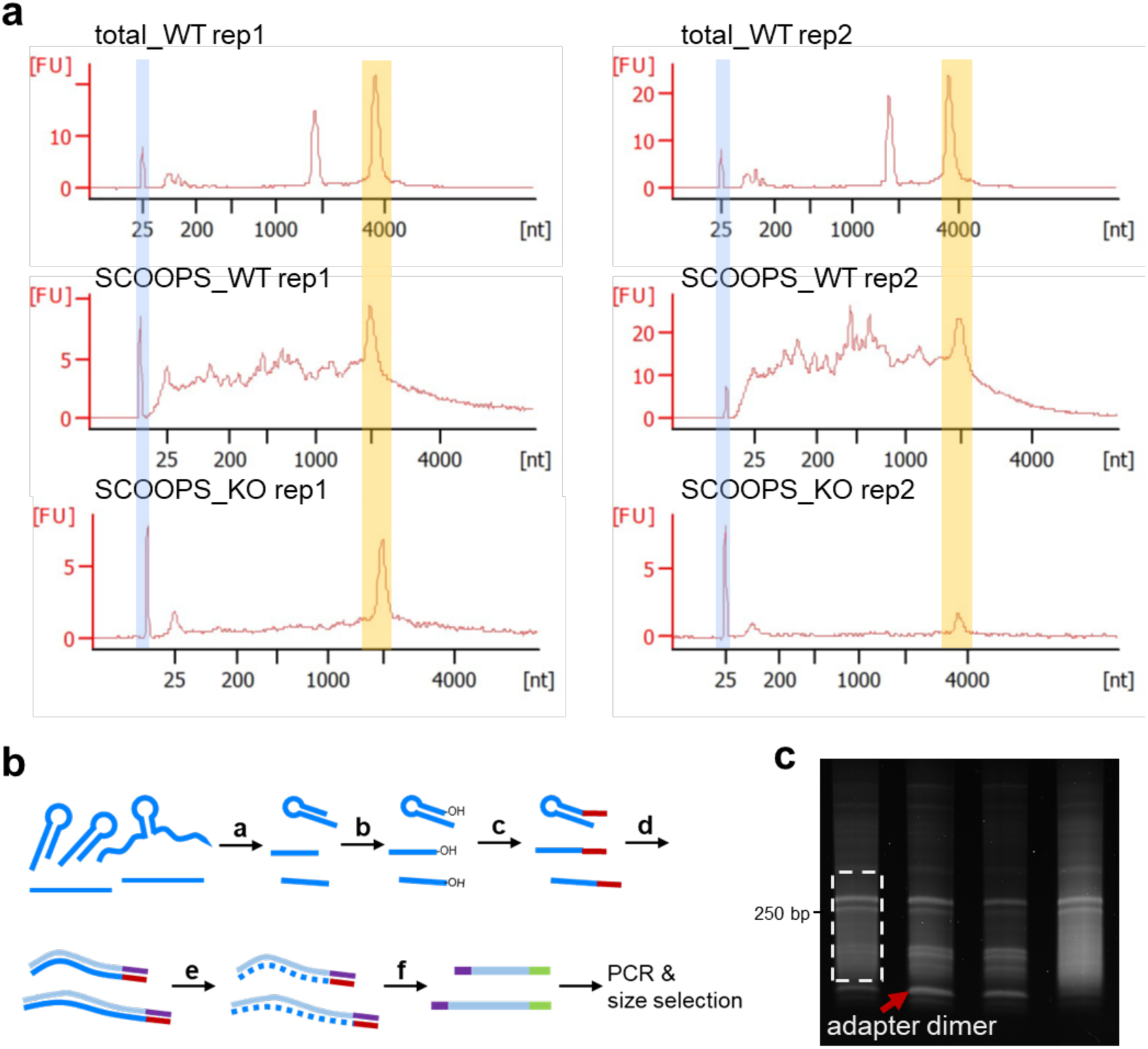
Supporting data for sequencing library preparation. **a**) Bioanalyzer electropherograms of all samples prior to RNA fragmentation. Blue shaded area indicates lower marker, and yellow shaded area indicates shift of 28S ribosomal RNA. For SCOOPS_WT rep1 and rep2, and SCOOPS_KO rep1, the processing software incorrectly assigned the ladder positions. **b**) Schematic representation of library preparation procedures for all samples. (a) Magnesium-based fragmentation; (b) 3’-end dephosphorylation; (c) pre-adenylated adapter ligation at RNA 3’-end; (d) reverse transcription; (e) RNA removal using sodium hydroxide; (f) second adapter ligation at 3’end of complementary DNA. **c**) A representative gel for PCR product size selection. Area circled by a white dashed frame right above the adapter dimer band (indicated by a red arrow) was excised in each lane. Samples from left to right, SCOOPS_WT rep1, SCOOPS_KO rep1, SCOOPS_KO rep2, SCOOPS_WT rep2.

**Supplementary Figure 9.**
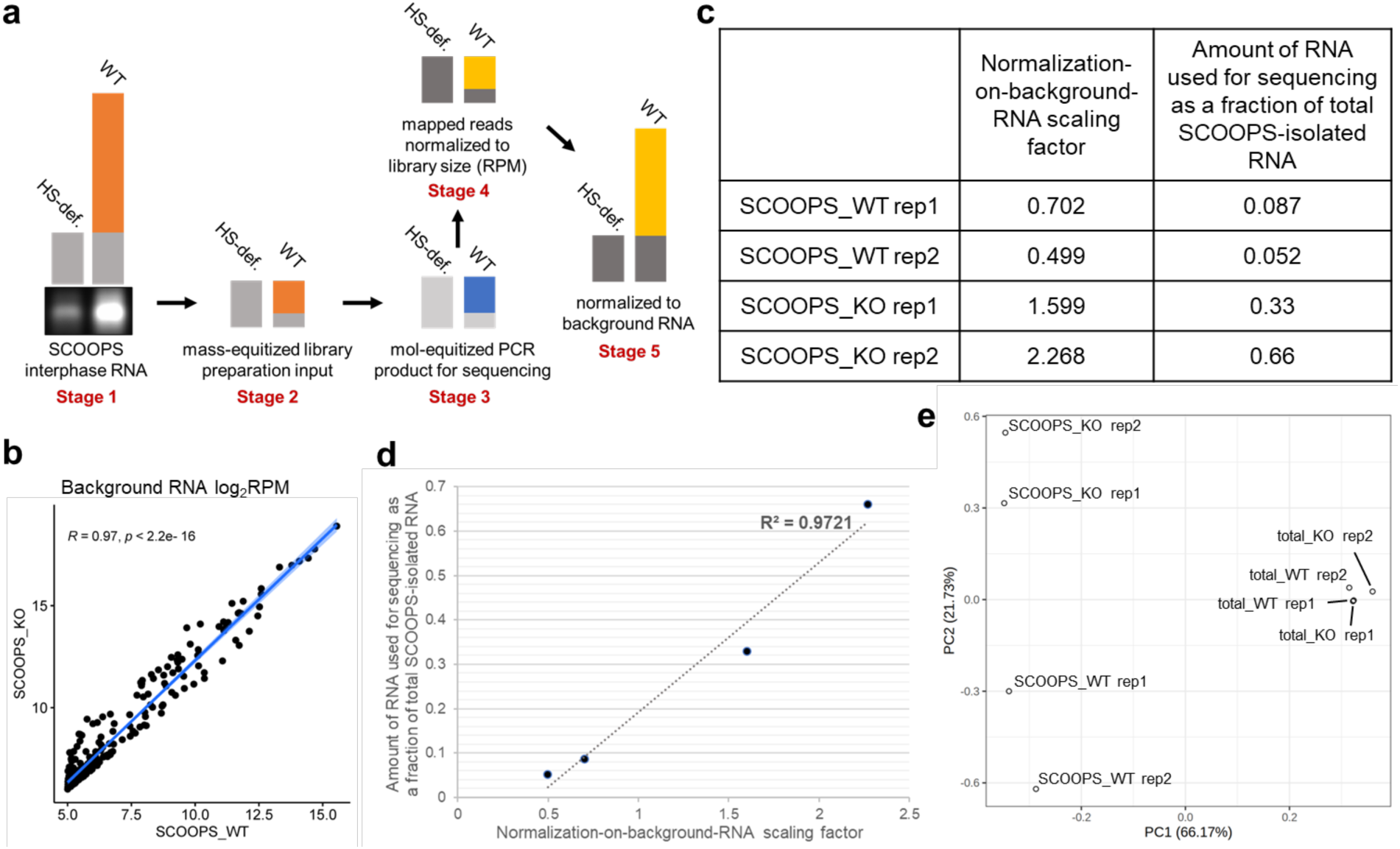
Additional data for RNA sequencing analysis. **a**) Illustration of rationales for normalization on background RNA prior to comparing SCOOPS_WT and KO samples. See **Supplementary Text 3** for a detailed description. HS-def., heparan sulfate-deficient sample, equivalent to EXT-KO. Grey bar stands for background RNA species in all stages. **b**) Pearson correlation analysis revealed background RNA species (see **Supplementary Text 3** for definition) from SCOOPS_WT and KO samples (prior to normalization on background RNA) were highly similar (R = 0.97). RPM of background RNA transcripts larger than 32 in both samples were log2-transformed prior to plotting the scatter plot. **c**) and **d**) Library-specific scaling factors generated by median of ratios normalization correlates well with the amount of library preparation input (150 ng) as fractions of total SCOOPS-isolated RNA. To scale read counts for each library, RPM values were divided by the scaling factor. **e**) Principal component analysis (PCA) of all samples. Normalized read counts were filtered (transcripts with read count < 10 were removed) and log2-transformed.

**Supplementary Figure 10.**
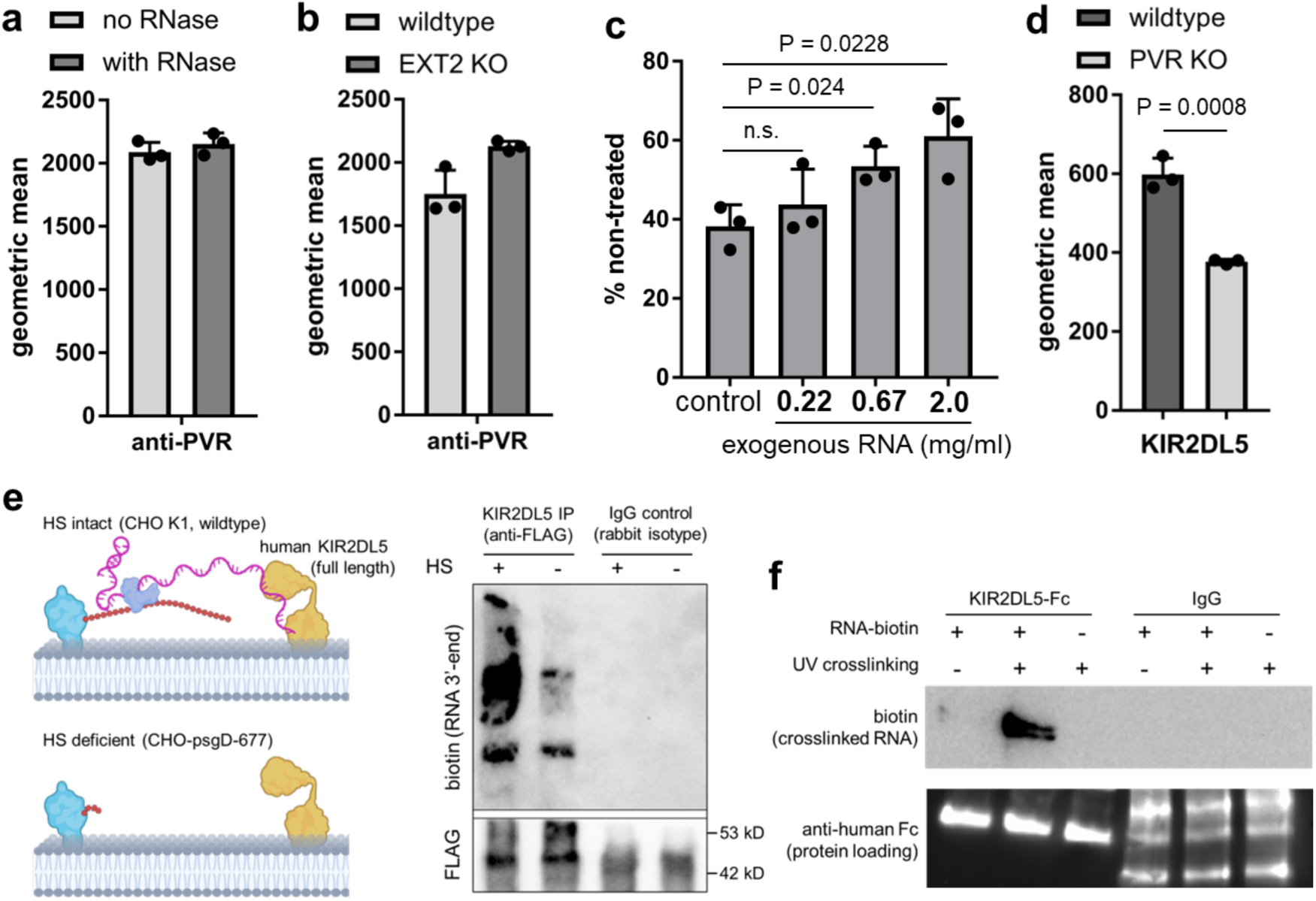
Supporting data for KIR2DL5 recruitment by hepRNA. **a**) Bar graphs from flow cytometry analysis of live cell-surface binding of monoclonal anti-PVR antibody, in the presence or absence of csRNA. Primary antibody was detected by AlexaFluor488-conjugated goat-anti-mouse secondary antibody. n = 3 independent cell cultures. **b**) Bar graphs from flow cytometry analysis of live cell-surface binding of monoclonal anti-PVR antibody to wildtype and EXT2-KO mutant Mel526 cells. n = 3 independent cell cultures. **c**) KIR2DL5-Fc binding was rescued by exogenous total RNA in a concentration-dependent manner. n = 3 independent cell cultures. **d**) Bar graphs from flow cytometry analysis of live cell surface binding of KIR2DL5-Fc to wildtype and PVR-KO mutant cells. n = 3 independent cell cultures. **e**) In-situ crosslinking of RNA to KIR2DL5 and immune-coprecipitation experiments in wildtype and HS-deficient cells. Precipitated KIR2DL5-bound RNA was on-bead labeled with pCp-biotin. FLAG tag was used to check for equal protein loading. Signals in IgG control were a result of polyclonal goat-anti-rabbit IgG HRP conjugate recognizing rabbit IgG released from beads. **f**) Western blot analysis of in-vitro crosslinked KIR2DL5-Fc and biotnylated RNA. Human IgG isotype control was included to check for non-specific binding.

**Supplementary Figure 11.**
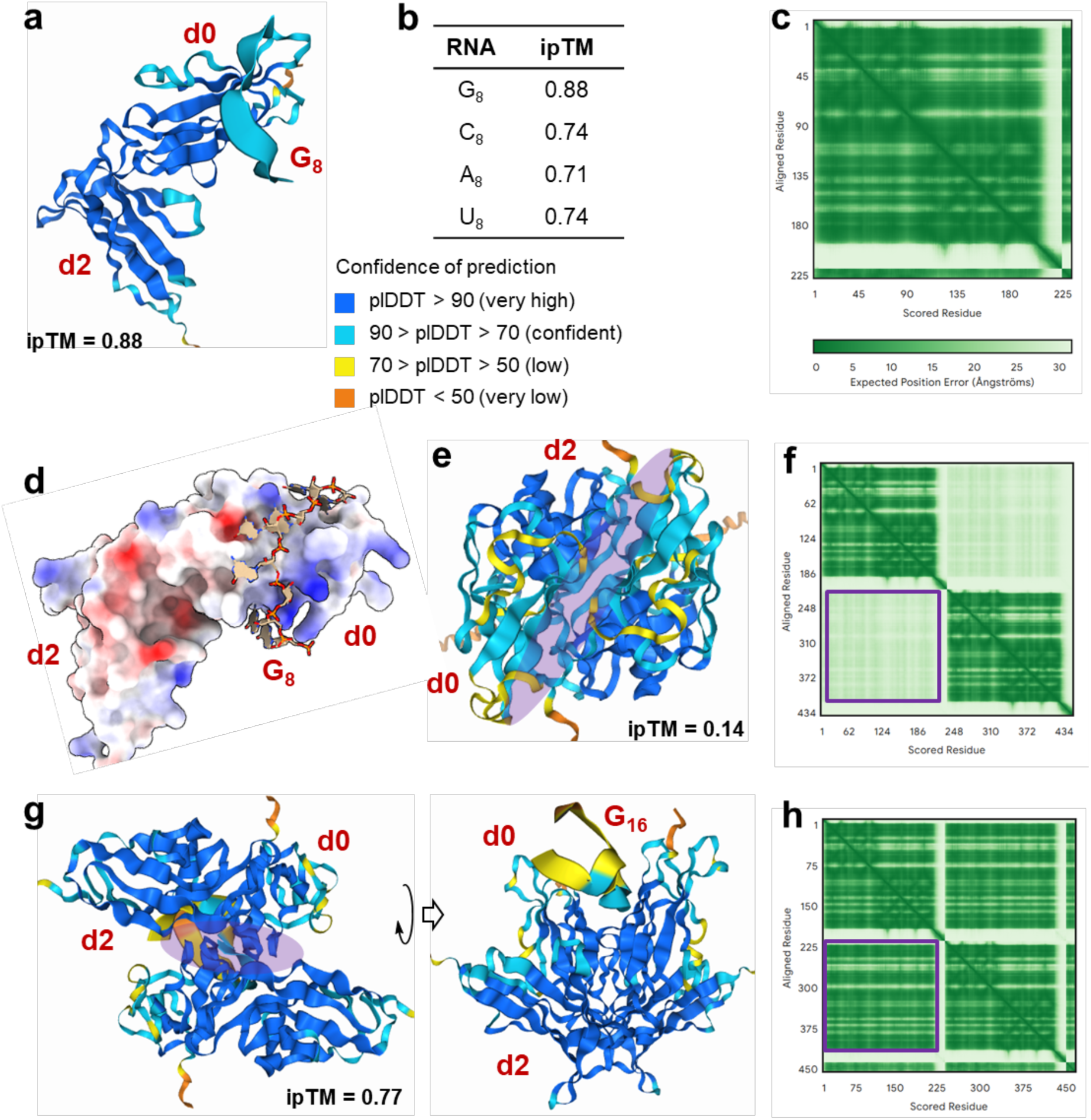
AlphaFold3-predicted structural model for KIR2DL5-RNA interaction. **a**) Predicted model for ectodomain of KIR2DL5 interacting with octaguanosine (G_8_) RNA. Immunoglobulin domains d0 and d2 of KIR2DL5 were indicated as red labels. The ribbon structures were colored by predicted local distance difference test (plDDT) scores, a measure of local confidence. See **Supplementary Table 8** for input sequences for AlphaFold3 prediction. **b**) Interface predicted template modeling (ipTM) scores for the KIR2DL5 ectodomain sequence and difference octanucleotide RNA oligos. Confident high-quality predictions are with values higher than 0.8. **c**) Predicted aligned error (PAE) matrix for KIR2DL5 ectodomain in complex with G_8_. **d**) Electrostatic surface potential of KIR2DL5 ectodomain. Immunoglobulin domain d0 contains a patch of positively charged residues proximal to the interphase between RNA and the protein. **e**) and **f**) Predicted model for dimerization of ectodomain of KIR2DL5 (protein only) and the corresponding predicted aligned error (PAE) matrix. The interphase between the two monomers is indicated by a purple shade. **g**) and **h**) Predicted model for dimerization of ectodomain of KIR2DL5 in the presence of a 16-mer RNA of guanosine (G_16_) and the corresponding predicted aligned error (PAE) matrix.

